# Multimodal layer-crossing interrogation of brain circuits enabled by microfluidic axialtrodes

**DOI:** 10.1101/2024.10.18.619110

**Authors:** Kunyang Sui, Neela K. Codadu, Daman Rathore, Guanghui Li, Marcello Meneghetti, Anders L. Bøcker, Rune W. Berg, Rob C. Wykes, Christos Markos

**Affiliations:** DTU Electro, Department of Electrical and Photonics Engineering, Technical University of Denmark, Kgs. Lyngby, 2800, Denmark; Department of Clinical and Experimental Epilepsy, Queen Square Institute of Neurology, University College London, London, WC1N 3BG, UK; Department of Neuroscience, Faculty of Health and Medical Sciences, University of Copenhagen, Copenhagen, 2200, Denmark; Centre for Nanotechnology in Medicine & Division of Neuroscience, Faculty of Biology Medicine & Health, University of Manchester, Manchester, UK; NORBLIS ApS, Virumgade 35D, 2830 Virum, Denmark

## Abstract

Modulating and recording neuronal activity are essential for probing brain function and developing therapies for neurological disorders. However, conventional flat-end optical fibers—widely used for deep brain access— interact with neural tissue only at their distal tip, limiting spatial resolution across brain layers. To overcome this constraint, we introduce the microfluidic axialtrode: a soft, multimodal neural interface created by controllably angle-cleaving a thermally drawn, multimaterial optical fiber integrated with multiple metal electrodes and microfluidic channels. We demonstrate *in vivo* that this design enables spatially distributed optogenetics, multisite electrophysiological recording, and targeted drug delivery along the fiber’s axis, allowing simultaneous interaction with multiple neuronal layers. The axial configuration increases the functional interface with brain tissue, while the soft polymer construction and reduced cross-sectional footprint significantly suppress the inflammatory response compared to conventional silica fibers. Integration with a 3D-printed scaffold, fabricated from FDA-approved biocompatible resin, provides mechanical stability and compatibility with standard experimental hardware. The monolithic integration of these features positions the axialtrode as a scalable and versatile platform for next-generation neural interfacing.

---

Understanding the brain’s complex functions and pathological mechanisms requires advanced neural interfaces capable of precise and multimodal interactions with neural circuits^1,2^. Current conventional approaches often require separate devices for optogenetics, drug infusion, and electrophysiological recording, complicating experimental setups and restricting real-time multimodal investigations^3,4^. Multifunctional neural interfaces typically rely on rigid probes or microfabricated platforms, while these devices have enabled important insights into brain activity, they often suffer from tissue damage and inflammatory responses in chronic *in vivo* investigation, which derive from the mechanical mismatch between the rigid probe and the soft neural tissue^5,6^.

During the last ten years, polymer-based multi-material optical fibers have emerged as a versatile platform for developing multifunctional brain implants^7–12^. Their superior flexibility results in minimal inflammatory responses compared to their rigid counterparts^13^. However, all the reported fiber-based neural implants so far rely on functional structures that are located at the distal end of the fiber, which imposes a limitation on exploring brain structures whose cellular populations are distributed in a dorso-ventral direction (vertically to the brain surface), such as medial prefrontal cortex^14^. Furthermore, depth-distinguish neural modulation recordings become imperative when the relevant neural circuits span a considerable depth range within the brain, such as investigating seizures across cortical and hippocampal regions^15^. Although the issue can be addressed by either positioning a flat-end facet optical fiber above electrode arrays or employing a tapered optical fiber beside electrode arrays^16^, the integration of a neural interface that combines multifunctionality is indispensable to mitigate relative positional uncertainties between sites of optical neuromodulation and neural activity recording at varying depths within the brain.

In this study, we introduce the microfluidic axialtrode (mAxialtrode) interface designed to overcome the existing limitations of the flat-end optical fibers. The functional structures, such as microfluidic channels (MCs) or electrodes, can be integrated into the cladding of a polymer step-index fiber. By precisely controlling an angled-cleaved end facet, we enable targeted drug delivery or neural recordings at varying positions along the fiber axial direction (distributed at different brain depths after implantation), as illustrated in Fig. 1a and b. *In vivo* experiments demonstrated the reliable performance of the proposed device, successfully achieving simultaneous optical stimulation and electrical signal recording across different brain structures (Fig. 1c). The “axial” drug delivery capability was validated by injecting two pharmaceutical compounds into different vertical locations across up to 2.7 mm deep in the brain tissue. This range allows targeted drug delivery spanning from the cortex to the thalamus in the rodent brains (Fig. 1d). Additionally, two kinds of 3D-printed interface scaffolds, fabricated using Formlabs Biomed Clear (a biocompatible, FDA-approved resin) and Phrozen resin, were developed to seamlessly connect the mAxialtrodes with external systems such as a light source, EEG recording system, and syringe pump (Fig. 1e,f). This innovative platform provides a versatile and minimally invasive tool for precise neuromodulation, offering new opportunities for advanced brain research and therapeutic interventions.

**Fig. 1:**
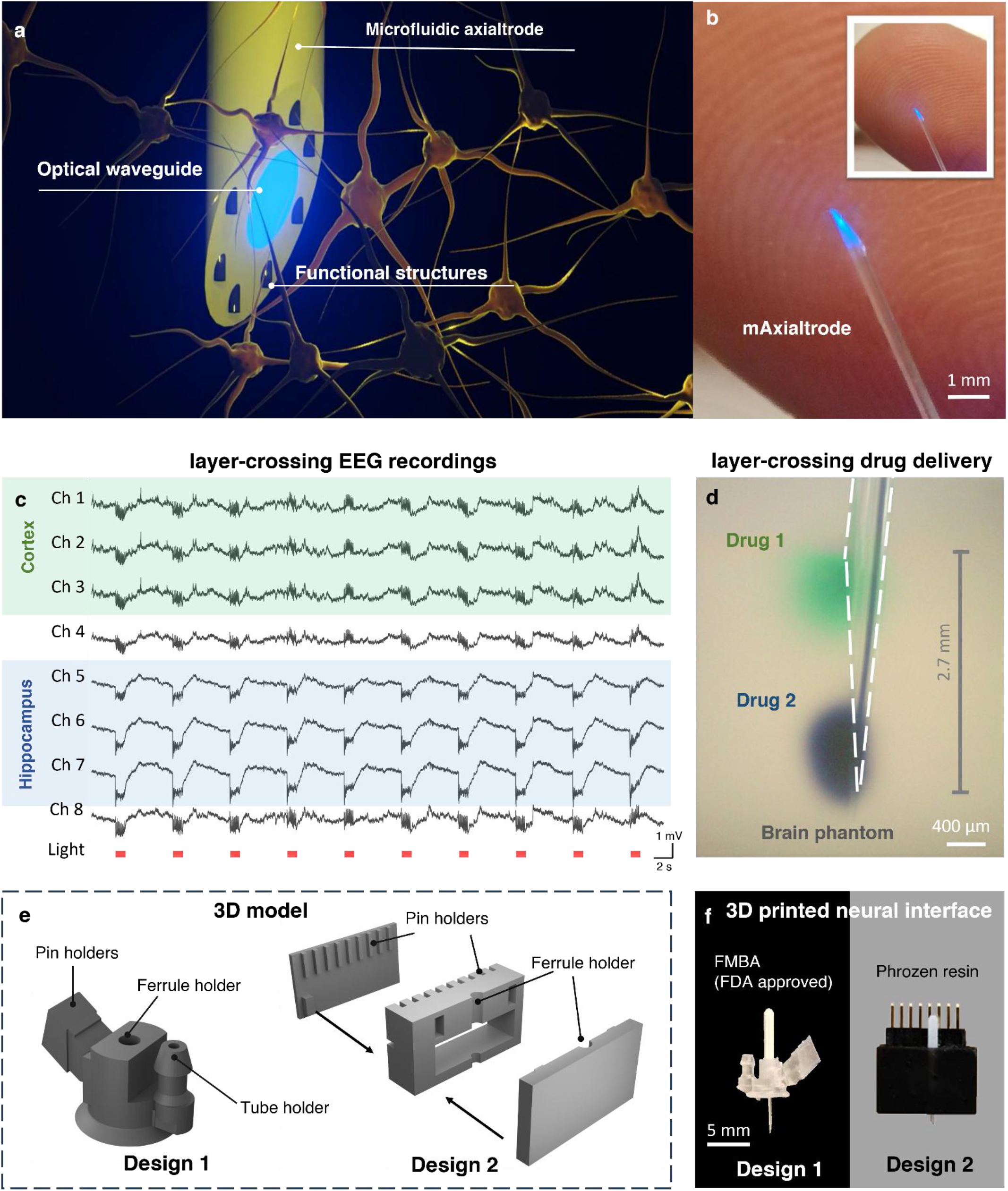
The mAxialtrode devices. **a**, Schematic of the mAxialtrode showing integrated optical waveguide, MCs, and electrodes for spatially distributed optogenetics, layer-crossing EEG recordings, and drug delivery. **b**, Photograph of the mAxialtrode tip placed on a fingertip for scale reference. **c**, EEG recordings from multiple channels spanning the cortex and hippocampus during light stimulation. **d**, Drug delivery at distinct depths in a brain phantom. Green dye was delivered from the uppermost MC and trypan blue from the lowest channel, demonstrating a delivery span of up to 2.7 mm. **e**, 3D models of two scaffold designs engineered to support and align the mAxialtrode. **f**, Image of the fully assembled device showing the mAxialtrode integrated with the 3D-printed biocompatible scaffold.

## Result

### Design and fabrication of mAxialtrode devices

The mAxialtrode is based on a step-index polymer optical fiber fabricated using a thermal drawing process. Specifically, polycarbonate (PC) and poly(methyl methacrylate) (PMMA) were chosen as the core and cladding materials due to their thermo-mechanical compatibility, overlapping viscosity-temperature profiles^17^, and distinct refractive indices^18^. During thermal drawing, a macroscopic preform, prepared by rod-in-tube method as depicted in Fig. 2a, was heated and drawn into a fiber with equivalent cross-sectional geometry but micro-scale dimensions. It is worth noting that the eight blind bores (hollow channels) with a diameter of 4 mm were added from the lower end-faceof the PMMA tube. These blind bores can help to build up constant pressure in the hollow channels to maintain the microstructure of the fibers during the thermal drawing process (Fig. 2b). In this study, the target fiber diameter ranged between 420 and 440 µm for compatibility with commercially available ceramic ferrules (CFLC440-10, Thorlabs). Several hundred meters of flexible polymer optical fiber were successfully produced in a single drawing (Supplementary Fig. 1). The cross-section of the drawn PC/PMMA fibers with a core diameter of 200 µm and microfluidic channel diameter of 20 µm is depicted in Fig. 2c, illustrating that the eight MCs are equally distributed around the fiber core in the PMMA. The fabricated optical fibers support light delivery at blue (470 nm wavelength) and red light (650 nm wavelength) for activating commonly used opsins in optogenetics, such as channelrhodopsin-2 (ChR2) and red-shifted rhodopsins, respectively ^19–21^ (Fig. 2c). The produced polymer optical fibers exhibit a low bend loss, as seen in Fig.2d, blue light can be guided even if the fiber is twisted around a small glass rod with a diameter of 5 mm. To achieve layer-crossing drug delivery, an angled-end facet was introduced to the polymer fibers to make the MCs distribute distinctive positions along the axial direction of the fibers. Exploiting the soft nature of polymer materials, we developed a controllable and accurate mechanical cleaving method for introducing the angled tip based on a custom stainless-steel mold (Supplementary Fig. 2). The mold features “V” grooves machined onto one surface of a stainless-steel block, with the angle between the grooves and the block’s edge customized according to the desired angle. During the cleaving process, the fiber was placed in one of the grooves with its tip extending out. Pressure was applied to the upper face of the mold to enhance friction between the fiber and the mold, preventing any unintended movement during cutting. Ultimately, the fiber tip was cleaved by sliding a blade along the side face of the stainless-steel block. Angled structures at 90° (flat-cleave), 45°, 30°, and 15° were produced based on this method, as shown in Fig. 2e. For electrophysiology (EEG) recordings, metal wires can be placed into the MCs and work as electrodes. As seen in Fig. 2f, the MCs within the cladding of the angled-tip polymer optical fiber were, mechanically exposed from the side surface of the fiber. Tungsten metal wires with a diameter of 20 µm were then integrated into the MCs from the exposed site under an optical microscope until they appeared at the angled surface of the fiber. Subsequently, the position of the tungsten wire was fixed by using cyanoacrylate adhesive. Here, the cyanoacrylate adhesive can not only fix the tungsten wire position but also seal one end of the MCs to build a constant pressure in the gap between the microchannel and the tungsten. This can stop any backflow of cerebrospinal fluid.

**Fig. 2:**
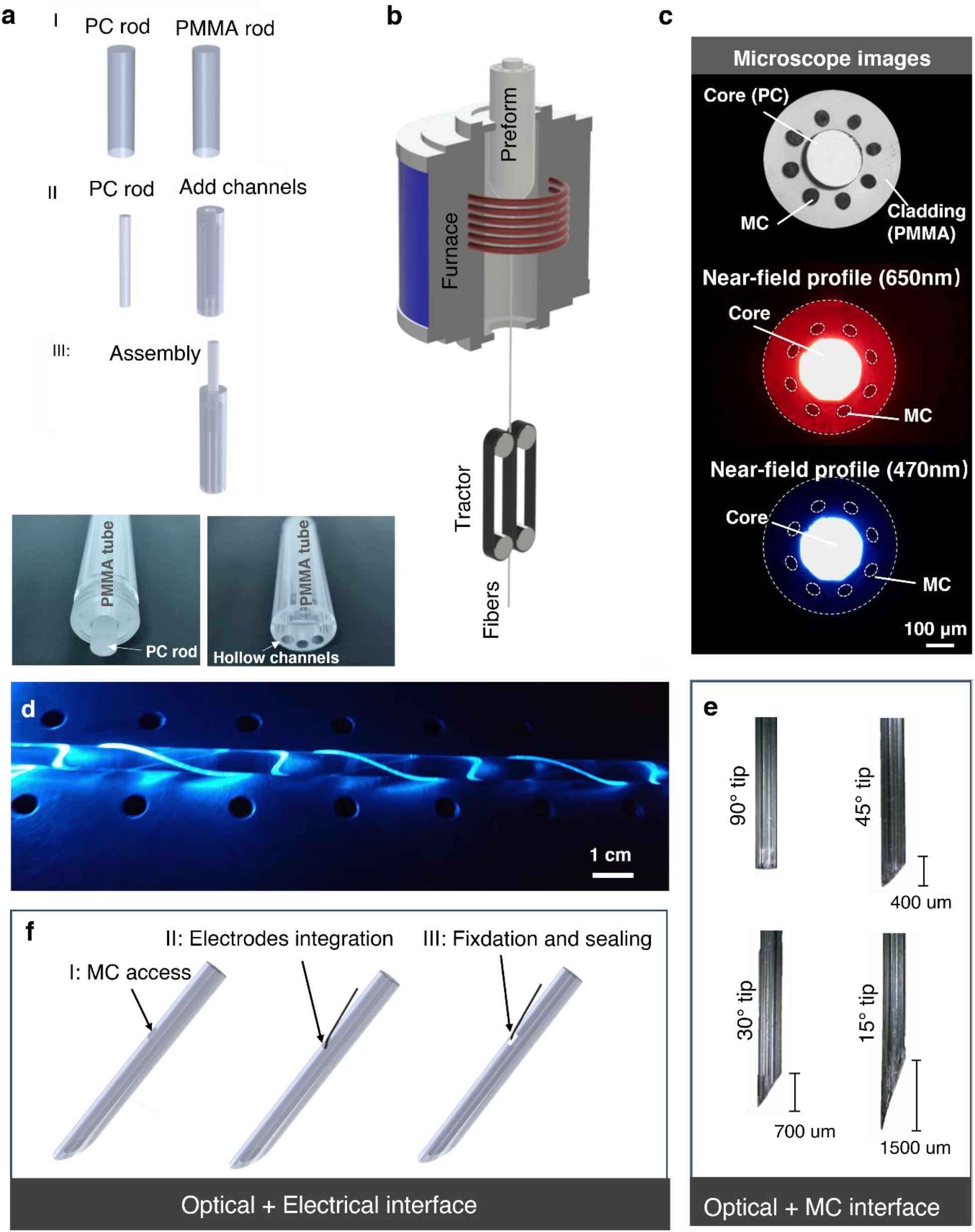
Development and fabrication of mAxialtrode. **a**, Preform fabrication process. I: Polycarbonate (PC) and poly(methyl methacrylate) (PMMA) rods were used as core and cladding materials, respectively. II: The PC rod was machined to a diameter of 10 mm, and a central hole (10.3 mm diameter) was drilled into the upper face of the PMMA rod to accommodate the PC core. Eight additional 4 mm-diameter holes were drilled into the lower face of the PMMA rod. III: The PC rod was inserted into the central channel of the PMMA rod to complete the preform. Bottom images show the assembled preform from both ends. **b**, Schematic of the thermal drawing process used to produce PC/PMMA optical fibers. Tens of meters of fiber were fabricated in a single-step fiber draw. **c**, Microscope image of the fiber cross-section showing the PC core surrounded by eight equidistant MCs (top). Near-field optical profiles for red (650 nm, middle) and blue (470 nm, bottom) light guided by the PC core. **d**, Image of the polymer optical fiber twisted around a 5 mm-diameter rod while maintaining blue light transmission through the core, demonstrating low bend loss. **e**, Optical microscopy images of fiber tips cleaved at different angles (90°, 45°, 30°, and 15°), showing the ability to control end-facet geometry for axial targeting. **f**, Electrode integration into the fiber. I: MCs were exposed from the fiber’s side surface using a scalpel. II: Tungsten wires (20 µm diameter) were inserted into the exposed MCs. III: Wires were fixed in place with a small drop of cyanoacrylate adhesive, which also sealed the channels to maintain pressure during *in vivo* use.

To facilitate the connection of the mAxialtrode with external equipment, including the light source for optical neuromodulation, EEG recording device, and drug delivery pump, custom scaffolds were created using a 3D printer (Sonic Mighty 8K, Phrozen). Here we proposed two scaffold designs (Fig.1 e f, and Supplementary Fig. 4): *Design 1* consists of a ferrule holder, pin holder, and a tube holder for the connection with an optical patch cable, electric pin header, and drug delivery tubes, respectively. To increase its biocompatibility in chronic research, the scaffold was printed with Formlabs Biomed Clear, an FDA-approved biocompatible resin. This scaffold was designed for simultaneous optical, electrical, and chemical neural modulation. Design 2 consisted of a ferrule holder for optogenetics and a pin holder with nine slots for multi-site neural activity recordings. This design can support up to 8 electrodes for EEG recordings across different depths. The multi-slot structures in design 2 are for preventing mutual entanglement of the metal wires or any potential unwanted cross-talking. Design 2 was printed using Phrozen resin for a cost-effective solution in acute experiments. The lightweight nature of the assembled mAxialtrode devices (0.25 g and 0.6 g for design 1 and 2, respectively) allows secure fixation on the heads of mice for *in vivo* experiments (Supplementary Video 1 and Supplementary Fig. 5).

### Characterization of the mAxialtrodes

The optical characterization of the mAxialtrodes was conducted by measuring their illumination maps and optical attenuation spectra. Illumination maps at the output end of the angled fiber were simulated using the ray-tracing software Zemax OpticStudio and experimentally verified in a brain slice. In the simulations, we compared the illumination map of angled-tip fibers (45° and 30°) with that of a flat-end optical fiber (90°) (Supplementary Fig. 6 for simulation details). The corresponding results are depicted in Fig. 3a. It can be seen that the medium adjacent to the end facet of the mAxialtrode was efficiently illuminated at different angles. Interestingly, with a reduction in the cleaving angle, part of the light propagates in the core of the fiber and is reflected by the interface between the core and the brain medium (forming *cluster b*) and further reflected by the interface between the cladding and the brain medium (forming *cluster c*) as illustrated in Supplementary Fig. 7. In addition, the decrease of the cleaving angle results in a reduced divergence of the emitted light (a smaller angle between the three clusters). Figure 3b compares the illumination map differences between fibers with flat end facets, 45°, and 30° angles investigated in real brain slices. It is evident that for the fiber with a 45° angled tip, part of the tissue on the back side of the end facet is illuminated due to reflection by the core-tissue interface, corresponding to the ray *cluster b* in the simulation (Supplementary Fig. 7). When the angle of the mAxialtrodes is reduced to 30°, more polymer volume must be inserted into the brain tissue for mapping the illumination profile due to the enlarged angled tip length. Despite the strongly compressed tissue around the fiber resulting in serious inhomogeneous tissue density and deformation of the illumination map, it is still evident that tissue near the tip of the fiber and on the back of the angled end facet has a high illumination intensity (Supplementary Fig. 8). To minimize tissue deformation during the experimental investigation of the illumination map, the orientation of the brain slice was rotated to maintain the upper surface of the slice in contact with the angled end facet of the 30° tip fiber in Fig. 3b.The attenuation spectrum of the optical waveguide in the wavelength range of 450 nm to 1100 nm was measured using the cut-back method^22–24^, as shown in Fig. 3c. The minimum loss was observed to be approximately ∼0.1 dB/cm between 750 nm and 850 nm, consistent with prior reports on PC-based polymer optical fibers^24^. High-loss bands are present between 600 nm and 1100 nm due to the high harmonics of carbon-hydrogen bonds’ vibration absorption^25^. The elevated loss region at wavelengths shorter than 600 nm is attributed to the scattering and electronic transition related to ultraviolet absorption^26^. Despite the pronounced loss in the blue region of the spectrum (470 nm), optical neuromodulation can still be achieved since only a short length of fiber is needed for implantation^11,12,24,27^.

**Fig. 3:**
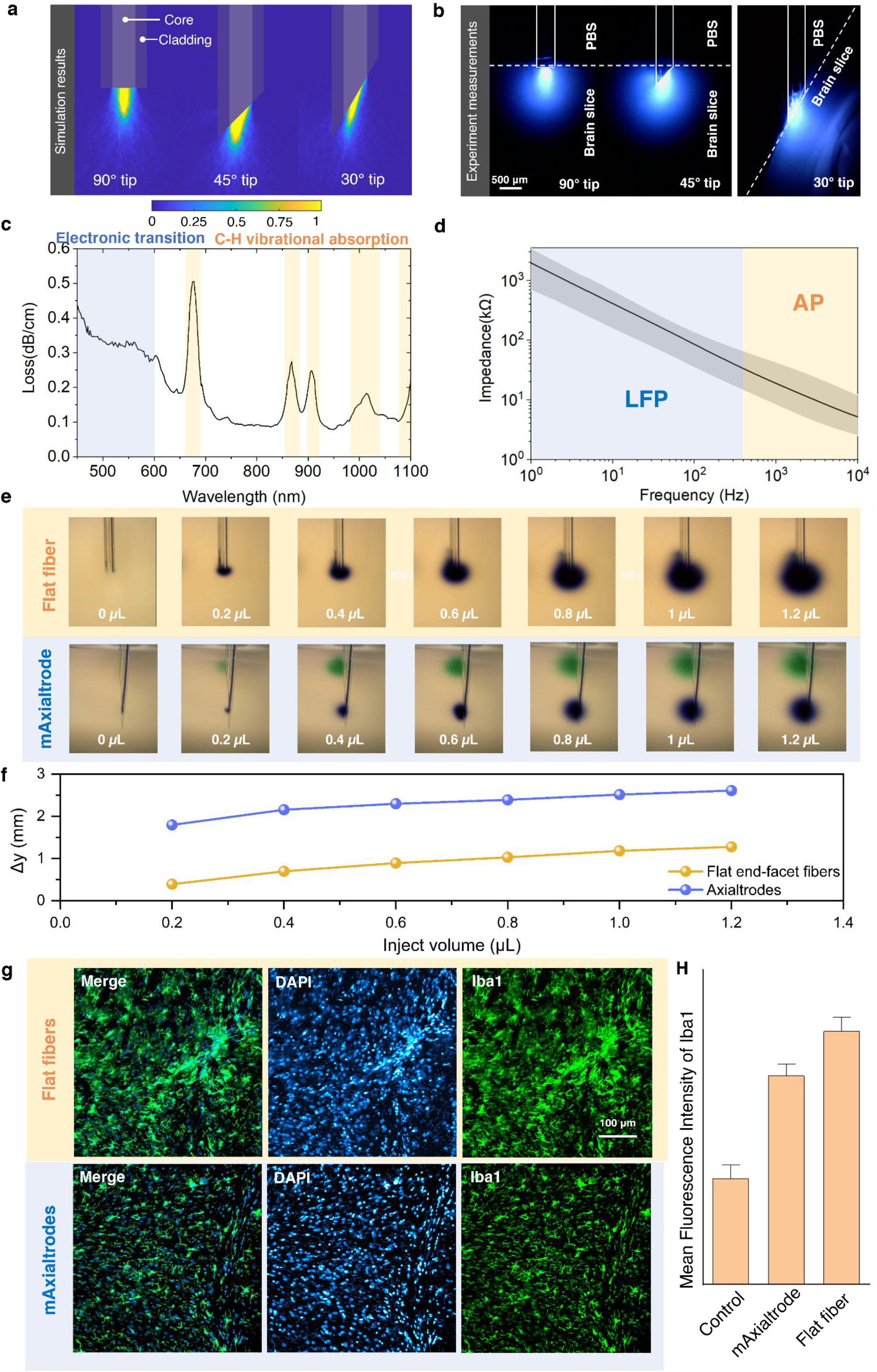
Characterization of the mAxialtrode. **a**, Simulated light distribution maps for fibers with tip angles of 90°, 45°, and 30°, illustrating changes in axial and radial illumination patterns. **b**, Experimental illumination maps of the same fiber tip geometries measured in fixed brain slices, confirming angle-dependent spatial light distribution. **c**, Loss profiles of polymer optical fibers, measured using the cut-back method using a supercontinuum source. **d**, Impedance spectra of integrated tungsten electrodes in the mAxialtrode across frequency. Data represents eight electrodes (n = 8); the black line shows the mean impedance, and the grey shaded area indicates variation across samples. **e**, Optical microscope images showing drug diffusion patterns from flat-cleaved and angled mAxialtrodes under equal injection volumes. **f**, Comparison of drug delivery depth span (Δy) between flat-end fibers and mAxialtrodes under equal total injection volume. For the mAxialtrode, half the volume was injected via the uppermost channel and the other half via the lowest, enabling axial separation of infusion zones. **g**, Immunohistochemistry images of microglia (Iba1, green) and nuclei (DAPI, blue) in brain tissue 4 weeks post-implantation of mAxialtrodes and conventional flat-end polymer fibers. **h**, Quantification of Iba1 fluorescence intensity in a 500 × 500 µm region centered at the fiber tip, comparing mAxialtrodes, flat-end fibers, and control (non-implanted) animals (n = 7 slices per group, 4-week implantation). Error bars represent standard error of the mean.

The electrical characterization of the neural interfaces involved measuring impedance spectroscopy with a chemical impedance analyzer (IM3590, Hioki). Fig. 3d displays the measured impedance of the eight electrodes in the frequency between 1 Hz and 10 kHz (more detailed characterization results of the impedance and phase of the eight integrated electrodes can be seen in Supplementary Fig. 9). Considering electrodes with impedances less than 1 MΩ at 1 kHz region are considered to be suitable for electrophysiology^28^, the integrated tungsten electrodes are anticipated to provide high-quality signal recordings, as we show in the subsequent *in vivo* experiments.

The drug delivery ability of the mAxialtrode was first characterized by measuring the build-up pressure (difference between the inner pressure in the tube and the atmospheric pressure) in channels under different drug delivery rates, since too high a build-up pressure may lead to tissue damage and device leakage or failure. From Supplementary Fig. 10a, the MCs have a negligible build-up pressure (<0.1 bar) under a delivery rate lower than 5 μL/min. This suggests that these MCs can achieve safe and efficient virus injection in optogenetic experiments, which require an injection rate lower than 1 μL/min^29,30^. This can be further confirmed by the output rate was found to be equal to the delivery rate when the delivery rate was set to be below 1 µL/min as seen in Supplementary Fig. 10b. The blowoff pressure (the pressure at which the tube expands too much and leads to the failure or leakage) has been measured to be ∼6 bar, which is higher than the necessary build-in pressure for convection-enhanced delivery^31^ (a technique used to deliver therapeutic agents, such as drugs, directly into the brain or spinal cord through a pressure-driven flow). To illustrate the depth-distinguished drug delivery function of the mAxialtrode, we compare its drug delivery pattern with that of conventional flat end-facet fiber, as seen in Fig. 3e. Two drugs with different colors, green and blue, were delivered by the uppermost channels and the lowest channels of the mAxialtrode, respectively. The drug delivery depth span can be up to 2.7 mm (Fig. 3f). In contrast, that of the flat-end facet fiber is 1.27 mm when the same volume of the drug was delivered, which is less than half of the depth span of the mAxialtrode.

The biocompatibility of the mAxialtrodes was investigated through immunohistochemical (IHC) evaluation of the microglia formation at the implantation sites. The inflammatory response differences between the mAxialtrode with a 15° angle tip and the conventional flat end-facet fiber with the same materials and diameter have been experimentally investigated. Here we chose the mAxialtrode with a tip angle of 15° since the long distance between its uppermost and lowest electrodes can be as large as 900 ± 10 µm, which is preferable for resolving the depth brain neural activity dynamics and recording in the following *in vivo* experiments. Mice with mAxialtrodes (N = 2) and standard end-facet fiber (N = 2) implanted were transcardially perfused 4 weeks after the implantation. The ones without any implantation surgeries served as control samples (N = 2). After the brains were dissected and sliced using a cryostat, the slices were stained with marker ionized calcium-binding adaptor molecule 1 (Iba1) and 4′,6-diamidino-2-phenylindole (DAPI) for activated microglial and their nuclei detection, respectively (Fig. 3g). The intensity of fluorescence observed within specified regions of interest (ROI) (500 µm x 500 µm) centered at the tip of the inserted fiber was used as an indicator of the reactive microglia induced in the brain by the neural devices. Quantitative analysis across different brain slices indicates a variance in the mean fluorescence intensity levels among the three distinct groups of samples (Fig. 3h), with the mAxialtrode proposed in this work producing less inflammatory response aggression compared with its flat end-facet counterpart (p = 0.0547, Supplementary Fig. 11). This can be attributed to the reduced foreign body response introduced by the smaller volume of the angled end-facet of the implant combined with the fact that the sharpened mAxialtrode’s tip reduces the tissue compression beneath the device during the implantation process compared with the fiber with a flat end-facet.

### In-vitro photoelectric artifacts evaluation

During electrophysiological neural activity recording, photoelectric artifacts represent a primary noise factor leading to data contamination and distortion^32^. For metallic materials, the valence and conduction bands typically overlap, making it easy for electrons to be excited by photon energy during light stimulation and result in photoelectric artifacts^33,34^. In addition, the photoelectric artifacts can also be affected by the distance between the optical waveguide and the electrodes^35^. Consequently, it is imperative to investigate the photoelectric artifacts of the proposed mAxialtrodes to ensure reliable signal recording.

With the assistance of the 3D printed scaffold and a custom-made pin adaptor, the proposed mAxialtrodes can be seamlessly connected with commercially available light sources and electrophysiology recording systems for all *in vitro* and *in vivo* experiments (Supplementary Fig. 12). *In vitro* measurements of optical stimulation-induced photoelectric artifacts were conducted by coupling a pulsed LED source (470 nm, 10 ms, 10 Hz) to the mAxialtrode with a 15° tip in phosphate-buffered saline. The position of each electrode channel number of the device can be seen in Fig. 4a, while Fig. 4b illustrates the temporal relationship between the stimulating light pulse and the resulting artifacts. The photoelectric signals are observed to be triggered at the rising edge of the laser pulse and begin to decrease at the falling edge. Recorded electrical signals from all eight electrodes during blue light stimulation in broadband (1-9000 Hz), the local field potential region (0.1-100 Hz), and the multi-unit activity region (500-9000 Hz) are presented in Fig. 4c (the full recordings are illustrated in Supplementary Fig. 13). Signals from the upper three electrodes are highlighted with a blue shadow, while signals from the lower three electrodes are marked with a red shadow. It is clear that the lower three electrodes at the angled end facet of the mAxialtrode experience light-induced electric artifacts, while the light stimulation has a negligible effect on the electrodes at other positions. A comparative analysis reveals that the electrode at the lowest position (connected with Ch6) exhibits the highest photoelectric artifact amplitude. This is attributed to the tendency of light propagating in the core of the fiber to emit at the tip of the fiber, as indicated by our simulations. The resulting higher light intensity at the tip leads to stronger photoelectric artifacts for electrodes at lower positions. Furthermore, it is observed that the photoelectric artifacts strongly influence the local field potential region (0.1-100 Hz) but have a negligible impact on multi-unit spike recordings (500-10000Hz). Power spectrum analysis of electric signal recording suggests that light-evoked artifacts mainly occur at frequencies of 10 Hz, 20 Hz, 30 Hz, and 40 Hz, matching the frequency of the stimulated light and its overtones (Supplementary Fig. 14). Using high-pass filtering, it has been previously reported as a proposed strategy for mitigating artifacts in neural spike data^36^. To further assess the separability of the artifacts from background noise, the signal-to-noise ratio (SNR)^37^ of the photoelectric artifacts under various high-pass filters (from 100 Hz to 1000 Hz) was calculated (Supplementary Fig. 15), indicating that a 500 Hz high-pass filter can effectively inhibit the photoelectric artifacts to an amplitude comparable to the background noise level in neural activity recording (more details of the SNR analysis can be seen in Methods section).

**Fig. 4:**
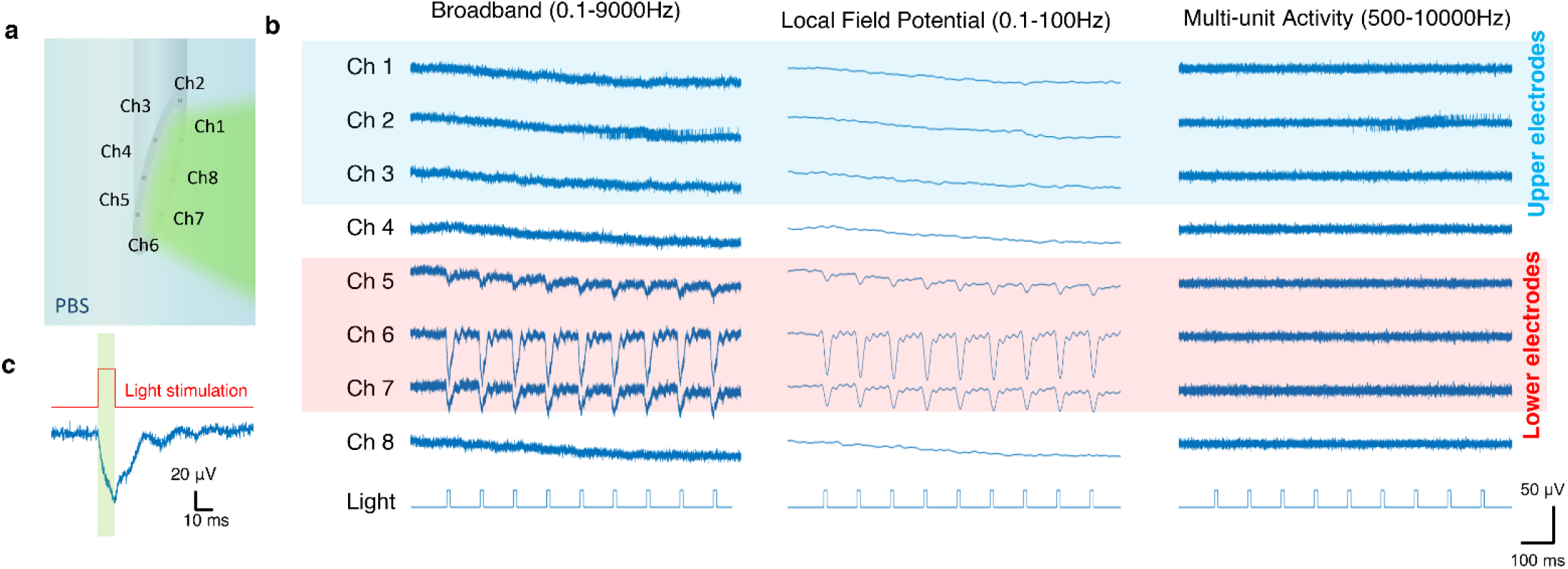
Characterization of the photoelectric artifacts of the mAxialtrode. **a**, Schematic representation of the channel number positions in the photoelectric artifacts characterization. **b**, Temporal relationship between the optical stimulation and the raised photoelectric artifact pulse. **c**, The recorded electric signal from all eight electrodes under illumination in broadband (1-9000 Hz), local field potential region (0.1-100 Hz), and multi-unit activity region (500-9000 Hz). The signals recorded from the upper three electrodes have been highlighted with a blue shadow, while the signals from the lower three electrodes have been marked with a red shadow.

### Awake in vivo theta wave recordings

Complex brain or spinal cord functions are underpinned by the synchronized activities of multiple neurons throughout different subregions of the brain. These neuronal ensembles are closely associated with various brain or spine rhythmic patterns^38–40^. Among these, theta frequency oscillations, predominantly ranging from 4 to 10 Hz, are a critical rhythm. Although theta frequency oscillations are detectable in both hippocampal and cortical regions^41^, it has been observed that the strength of cortical theta oscillations is typically less pronounced when compared to those in the hippocampus^42^.

In this study, we conducted a proof-of-concept experiment to assess the recording capabilities of the mAxialtrodes across different brain domains by recording the theta oscillation intensity difference between the cortex and the hippocampus regions. The concept of the experiment is illustrated in Fig. 5a. We implanted the 15°-based mAxialtrode into the brain of a mouse, positioning the bottom three electrodes within the hippocampal region and the top three electrodes in the cortical region. The span from the highest to the lowest electrode was measured to be ∼900 µm with the possibility of exceeding the 1 mm. A representative 10-second segment of the recordings from all eight electrodes is presented in Fig. 5b (full recordings can be found in Supplementary Fig. 16), revealing a high degree of waveform similarity across the electrodes. Yet, when a band-pass filter (4-10 Hz) was applied to isolate theta oscillations, a notable difference emerged: the amplitudes of theta rhythms recorded by the bottom three electrodes (hippocampus) were consistently higher than those captured by the top three electrodes (cortex), as demonstrated in Supplementary Fig. 17. This observation was further supported by power spectrum analysis of the recordings from each electrode, illustrated in Fig. 5c. Here, the theta rhythms recorded by the hippocampus-located electrodes displayed markedly higher power compared to those recorded by the cortex-located electrodes. This correlation between our experimental findings and the established literature^42^ validates the utility of our mAxialtrode for depth-specific neural activity recording. After the experiment, the implantation location was verified through an examination of the acute insertion footprint (the device was coated with Hoechst dye to serve as a position marker under microscopic examination before implantation). Figure 5d reveals the fluorescent imaging of the tissue damage caused by the mAxialtrode implantation in dorsal-ventral direction of the mouse brain. It can be seen that the insertion footprint penetrated through the cortical layer and extended into the hippocampal region of the brain.

**Fig. 5:**
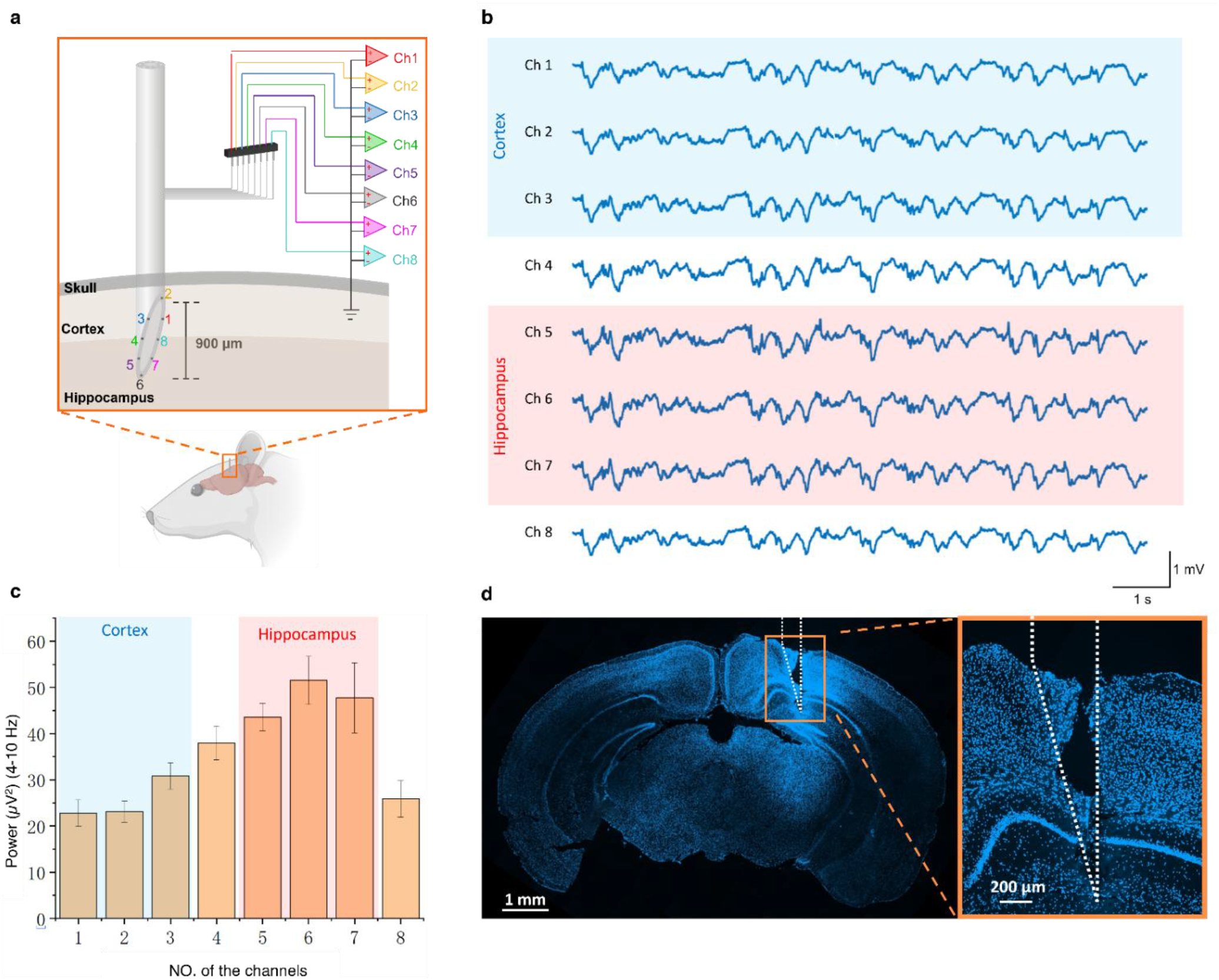
Distinguishing the theta rhythms in the cortex and hippocampus using the developed mAxialtrode. **a**, Illustration of the experimental concept: the mAxialtrode was implanted into the brain of a mouse with the lower three electrodes reaching the hippocampus region and the upper three electrodes touching the cortex tissue. **b**, The electric signals were recorded by the eight electrodes within 10 seconds. **c**, Power of recorded theta wave by all the eight channels of the mAxialtrodes (electrodes location at the cortex and hippocampus were highlighted with light blue and red shadows, respectively). **d**, Confocal fluorescent imaging of the insertion footprint in the dorsal-ventral direction of the mouse brain (magnification 10X).

### In vivo axial optogenetics and electrophysiology

Optogenetics stands out for its exceptional cell-type specificity, providing a powerful means to explore functional connectivity across various brain regions with precise spatial and neuronal population selectivity^43^. The integrated optical waveguide and electrodes in the mAxialtrodes make it an efficient tool for optogenetic applications. A proof-of-concept experiment was conducted to prove its utility in concurrent optical neural activity stimulation and electrophysiology recordings.

Supplementary Video 2 shows an animated rendition of the process of neural activity recording during an optogenetics experiment with the proposed mAxialtrode. After ChR2 was expressed in the visual cortex of a mouse brain, a mAxialtrode with a tip angle of 15° was loaded into the brain until the uppermost electrodes (which correspond to channel 2 in the schematic in Fig. 4a) reached the surface of the visual cortex. Neural activities were stimulated by delivering blue light from a fiber-coupled LED. The output power at the tip of the fiber was measured to be 1.4 mW. The frequency and the pulse width were modulated by an LED driver (frequency: 10 Hz, pulse width: 10 ms), and TTL pulses from the stimulating signal were synchronized with the electrophysiology recording via the Intan RHX program. The resultant signals recorded by the eight electrodes are presented in Supplementary Fig. 18 and 19. A high-pass filter (500 Hz) was applied to highlight the light-evoked neural activity, demonstrating that spikes triggered by optical stimulation can be recorded by all integrated electrodes (Supplementary Fig. 20). From a typical 1-second recording illustrated in Fig. 6a, the signal from electrode 6 (Ch6) exhibited a lower amplitude than the rest possibly due to its further position away from the opsins (the virus was injected into the visual cortex). Interestingly, electrodes at upper positions (Ch1 to Ch3) captured typical tri-phasic axonal signals, while electrodes at lower positions (Ch6 and Ch7) recorded multi-unit activities with more complex ‘w’ shapes. Among the 6 channels that can record single-unit neural activities (Ch1-Ch5 and Ch8), we use principal component analysis on Ch1 (a representative single-unit activity recording channel) and found that two distinct spike clusters were identified, as shown in Fig. 6b while Fig. 6c displays the superimposition and average shapes of these two spike types. The spike firing rate (SFR) was calculated by convolving the spikes’ temporal locations with a Gaussian kernel (bin size: 0.2 s) as shown in Fig. 6d. The distinct SFRs of the two spike units (Fig. 6e) make it obvious that one of the clusters is strongly related to the light stimulation (cluster 2 in Fig. 6c), while the other appeared randomly throughout the recording (cluster 1 in Fig. 6c).

**Fig. 6:**
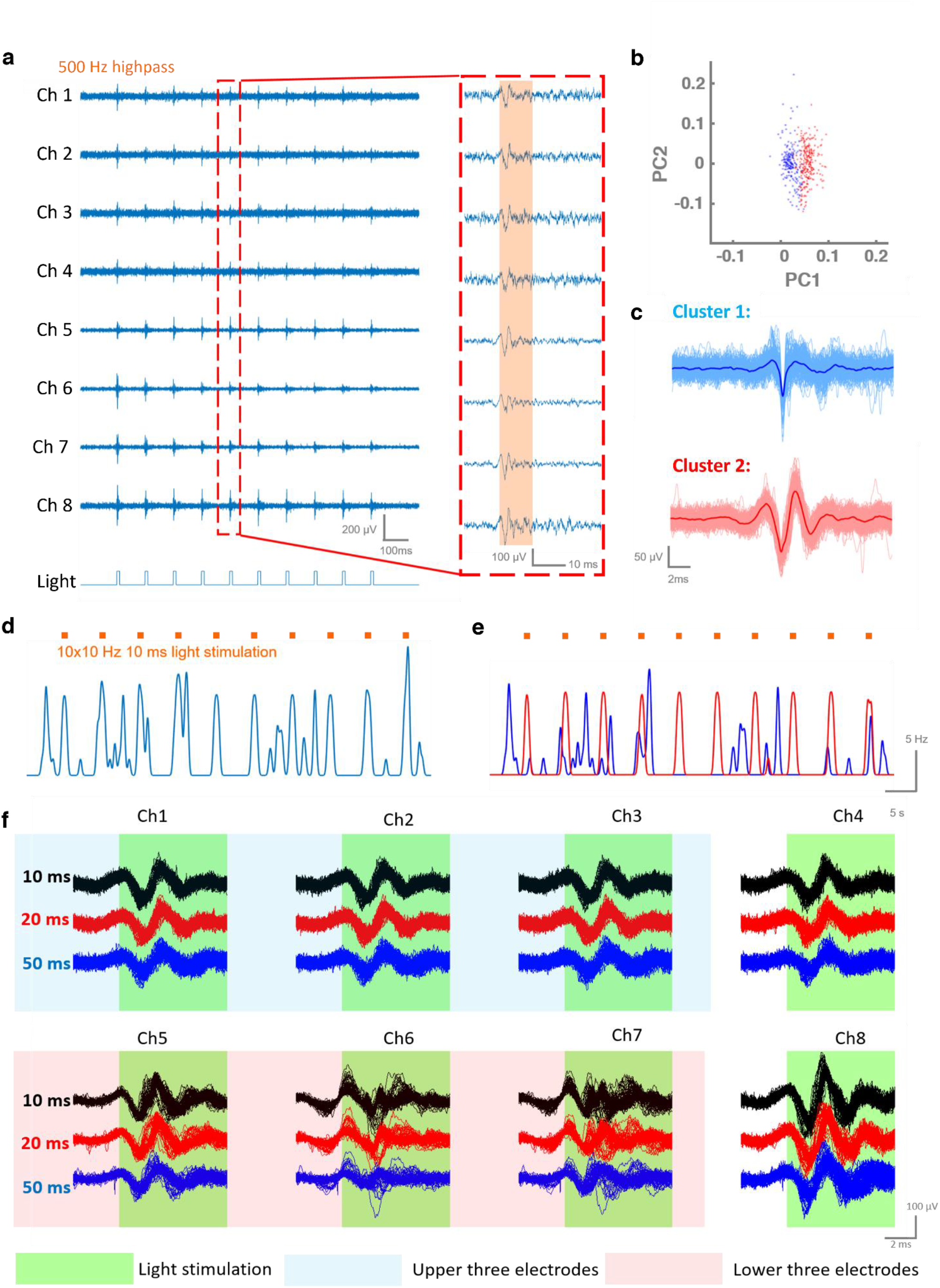
Optogenetic stimulation and electrophysiological recording using the mAxialtrode. **a**, Neural activity recorded simultaneously from eight integrated electrodes *in vivo*. **b**, Principal component analysis of spike data collected from Ch1, identifying two distinct units. **c**, Overlay and average waveforms of the two spike clusters identified in **b**, showing consistent action potential shapes. **d**, Spike firing rate recorded by Ch1 during blue light stimulation. **e**, Spike firing rates of the two PCA-sorted units under blue light stimulation, demonstrating cluster-specific modulation. **f**, Comparison of neural activity waveforms recorded across all eight electrodes during light stimulation with different pulse widths (10 ms, 20 ms, 50 ms), showing pulse-dependent response characteristics.

The study also investigated the effect of varying light pulse widths (10 ms, 20 ms, and 50 ms) on the evoked spikes. To do so, the spikes recorded by the eight channels immediately following light stimulation were compared (Fig. 6f). It was observed that a 10 ms light pulse evoked spikes with the highest amplitude, and the average amplitude of spikes collected by the eight electrodes decreased from 101.55 µV to 76.71 µV as the pulse width increased from 10 ms to 50 ms (Supplementary Fig. 21 and Table 2). The results can be explained through the kinetic model of the ChR2 photocycle, which involves four kinetic intermediate states (P1, P2, P3, and P4)^44^. It is hypothesized that the increase in photocurrents within the first 10 ms following light stimulation is attributed to the state transition from P1 to P3, where the open state time constants for transitions from P1 to P2 and P2 to P3 are 1 ms and 10 ms, respectively. However, when the duration of the light pulse exceeds 10 ms and reaches the P4 state, the desensitization leads to a reduction in photocurrents due to the inactivation of ChR2^45^.

## Discussion

We introduced and developed an implantable neural interface for layer-crossing interrogation of brain circuits. By integrating the proposed mAxialtrode with a 3D-printed biocompatible scaffold, our platform enables simultaneous optical stimulation, electrophysiological recording, and targeted drug delivery along the dorso-ventral (vertical) direction of the brain.

A key advantage of this interface lies in its structural versatility where critical device parameters such as implant diameter, optical waveguide dimensions, and the spatial arrangement of functional components can be readily tailored to specific experimental needs. The thermal drawing process allows for scalable fabrication of optical waveguides to match standard patch cable cores, ensuring efficient light coupling. Additionally, by precisely adjusting the cleaving angle of the fiber tip, the axial distribution of electrodes and microfluidic outlets can be customized to target distinct neuronal layers or brain regions.

Compared with flat-end facet fiber-based implants, the mAxialtrode can not only reduce the tissue compression during the implantation but also avoid buckling failure to increase the successful implantation rate^46,47^. This is extremely important for the development of chronic, minimally invasive, and reliable implants based on compliant materials (such as soft polymers)^48^. Furthermore, another important advantage of the mAxialtrode is its reduced volume compared to flat-end fibers, which eventually leads to a suppressed inflammatory response in chronic implantations.

The capability of the proposed mAxialtrodes in terms of depth-resolved neural activity recording has been demonstrated through the differentiation of theta rhythms in the cortical and hippocampal regions of the brain. The theta wave’s power, as recorded by the upper three electrodes of the mAxialtrode situated in the cortical region, has a lower intensity compared to that recorded by the lower three electrodes in the hippocampal region. During optogenetics experiment, neural activities were recorded by all eight electrodes of the fiber-based implant, triggered by pulse light from an LED for ChR2 stimulation. Although preliminary *in vitro* experiments indicated potential photoelectric artifacts in the low-frequency range on the three electrodes nearest the optical waveguide’s tip, these artifacts could be effectively eliminated by employing a high-pass filter during the action potential recordings.

Compared to the existing tapered fibers interfaces^49^, the mAxialtrode proposed in this study enables not only multiple brain region crossing EEG recordings but also target drug delivery, all without the need to reposition the fiber interface. Furthermore, this work explores critical aspects of the angled optical waveguide used for neuromodulation, including comparisons of illumination profiles between flat and different angled end facet fibers, as well as investigations into photoelectric artifacts associated with the angled optical waveguide during optogenetics. Details of the angled end facet introducing method have been described, along with an analysis of how tip angles influence the functional structure span along the fiber’s length.

In summary, mAxialtrodes overcome the limitation of single-layer brain neural modulation and recording posed by conventional flat-end optical fibers. Their extended axial distribution of functional components enables interrogation of neural dynamics across multiple brain layers or even controlled photo-pharmacology. The scaffold made from FDA-approved biocompatible resin enhances suitability for chronic implantation and ensures compatibility with standard experimental setups. Future developments could include wireless control and real-time sensing, advancing the device toward closed-loop neuromodulation and clinical neurotherapy applications.

## Methods

### Thermal drawing of the polymer optical fibers

PC and PMMA polymer rods with a diameter of 25 mm were acquired from Goodfellow UK. The diameter of the PC rod was reduced to 10 mm by machining and a central hole with a diameter of 10.3 mm was drilled in the PMMA rod. Subsequently, eight hollow channels, each 4 mm in diameter, were drilled at the alternate end of the PMMA rod using a precision drill bit (HSCO, EJOT). This assembly, with the PC rod inserted into the PMMA tube, served as the structured preform for fiber fabrication. The fibers were drawn using an optical fiber draw tower facility at DTU Electro. The preform was fixed to the feeding system using a custom metal adaptor and then loaded into the furnace. The drawing process started after the temperature of the furnace reached 200 °C, causing the preform to form a drop. The speed of the feed system was set to 0.2 mm/min, while the fiber drawing speed was adjusted based on the target fiber diameter. Throughout the fiber fabrication, a laser diameter gauge was utilized to continuously monitor and ensure the stability of the fiber’s diameter, which exhibited minimal fluctuation deviation (<1%) during several hundreds of meters.

### Microscope image and near-field optical profiles

The PC/PMMA polymer optical fiber was cleaved using stainless steel blades (EHDIS) and the cross-section images of the fibers were captured in the transmission mode of a Zeiss Axioscan A1 microscope, equipped with an Axiocam 305 color camera. To image the output light profile emanating from the PC/PMMA polymer optical fibers, a customized cage system was built to facilitate light coupling. Light emitted from a blue LED (M470L4, Thorlabs) or a red LED (M660L4, Thorlabs) was initially collimated through an aspheric lens (AL2520M-A, Thorlabs). The path of this collimated light was then adjusted using two broadband dielectric mirrors (BB1-E02, Thorlabs), each mounted on right-angle kinematic mounts (KCB1C/M, Thorlabs). The coupling into the fiber was achieved with another aspheric lens (AL2520M-A). The propagation light in the core of PC/PMMA was imaged by a digital microscope (AM7915MZT, Dino-Lite).

### Development of the mAxialtrode scaffold

The 3D model of the scaffold was designed using SolidWorks and subsequently fabricated using a high-resolution 3D printer (Sonic Mighty 8K, Phrozen), capable of achieving a resolution of 28 µm in the x-y plane (length and width) and 10 µm in the z-axis (thickness) with masked stereolithography apparatus (MSLA) technique. In the printing process, the build plate and resin vat were first installed in the printer. The Phrozen resin water was thoroughly mixed and then poured into the resin tank. The 3D model print file was transferred to the 3D printer via a USB disk to initiate the printing process. To minimize exposure to excessive UV light, the printer’s plastic case was kept closed during the print. The key printing parameters have been optimized as shown in Supplementary Fig. 3 and Table 1. Upon completion of printing, the build plate was removed from the printer. A metal scraper was then used to carefully detach the printed scaffolds from the build plate. The scaffolds were subsequently cleaned using Phrozen Wash Resin Cleaner and underwent final UV curing in a post-curing chamber. Remarkably, this printing process allows for the production of up to 30 scaffolds for design 1 and 16 scaffolds for design 2 within one hour.

### Ray tracing simulations and optical fiber illumination map in brain tissue

The simulation of light distribution for optical fibers with varying tip angles (90°, 45°, and 30°) was conducted using the ray-tracing software Zemax-OpticStudio. The layout of the model can be seen in Supplementary Fig. 6: the model consisted of two concentric cylinders representing a step-index optical fiber, with diameters of 200 µm and 400 µm, respectively. The refractive indices for the inner and outer cylinders were set at 1.58^50^ and 1.49^51^, corresponding to the properties of PC and PMMA materials. As ferrule patch cables with a numerical aperture of 0.39 (MR83L01, Thorlabs)^52,53^ are commonly used to couple the light from a light source to the optical waveguide in the probes, accordingly, a point source with a divergence angle of 23° (matching the light emission angle of a patch cable with an NA of 0.39) and a wavelength of 470 nm was positioned adjacent to one end of the fiber in the simulation. Additionally, a homogenous rectangular volume, representing brain tissue, was placed at the opposite end of the fiber. The refractive index of the brain tissue volume was set as 1.36 and its scattering properties were modeled using a Henyey-Greenstein scattering function with a mean free path of 0.04895 mm, an anisotropy value of 0.9254, and a transmission of 0.9989^54^. The spatial power distribution at the end of the fiber was recorded by raster scanning a square detector (dimension: 10×10 μm) in the x-z plate. The resulting light intensity distribution was mapped in MATLAB. An interpolation algorithm was applied to enhance the resolution of the resultant image.

### Fiber loss measurement

The transmission properties of the fabricated fibers were characterized by measuring the propagation loss using the standard cut-back method. A high-power supercontinuum fiber laser (SuperK Extreme, NKT Photonics A/S) covering a wide wavelength range from 400 nm to 2400 nm was utilized as the light source. Due to the necessity of high pump power for generating broadband supercontinuum light, resulting in a total average power exceeding 5 W, a variable neutral density filter at the output of the laser was employed to reduce the launch power to ∼100 mW, thereby preventing potential thermal damage to the end facets of the fiber. In addition, a beam trap (BT610/M, Thorlabs) was used to safely capture any residual reflected light. The light from the laser source was then efficiently coupled into the core of the polymer optical fibers using an achromatic objective lens with a 10x magnification and the output transmitted spectrum was measured using a spectrometer (CCS200, Thorlabs).

### Impedance spectroscopy measurements

For the electrochemical impedance spectroscopy analysis, a chemical impedance analyzer (IM3590, Hioki) was employed in a three-terminal setup. In this configuration, a tungsten microwire, coiled around a rod, functioned as the counter electrode. An Ag/AgCl electrode was chosen as the reference electrode, and the electrodes integrated into the angled tip fiber devices served as the working electrodes. All electrodes were immersed in a phosphate-buffered saline (PBS) solution, which acted as the electrolytic medium. The impedance properties of the fiber devices were then meticulously measured across a frequency range spanning from 1 Hz to 10 kHz.

### Setup for the pressure measurement in the MCs during drug delivery

The pressure within the drug delivery tube was monitored using a Wika A-10 pressure gauge, which outputs a current proportional to the applied pressure. A 250 Ω resistor was employed to convert the current to a voltage signal, which was then measured by an Analog Discovery 2 digital oscilloscope with a resolution of 0.005 bar. For liquid pressure testing, the Wika A-10 was mounted onto the upper face of a custom adapter device, designed and fabricated by DTU Space Mechanics. One side of the adaptor was connected to a syringe pump (Harvard Apparatus, PHD 2000), while the other side was linked to the microfluidic axiotrode. As the syringe pump generated pressure during drug delivery, the inner chamber of the adaptor experienced the same pressure as the MC,s which can be recorded by the integrated pressure gauge.

### Electrophysiology recordings

Electrophysiological data acquisition was conducted using the RHS Stim/Recording System from Intan Technologies, which comprises an RHS Stim/Recording Headstage, an RHS Stim/Recording Controller, and a Stim SPI Interface Cable. The electrophysiology setup is detailed in Supplementary Fig. 11. A self-made adaptor was developed for delivering the electrical signals from eight pins in the neural interface device to the RHS stim/recording head stage. All the results can be visualized by the RHX Data Acquisition Software during the *in vivo* experiment. For all the recordings, the sampling rate was set at 20 kHz.

### Optical Stimulation

Pulsed light at the wavelength of 470 nm (10Hz) with various pulse widths for optical stimulation was generated from a fiber-coupled LED (M470L4, Thorlabs) and driven by an LED driver (DC2200, Thorlabs) with internal modulation of square waveforms. The power output at the fiber tip was calibrated to 1.4 mW. For synchronizing the temporal profiles of optical stimulation pulses with the electrical signals, the LED driver was integrated with the Intan data acquisition system, enabling cohesive data communication. Identical optical stimulation parameters were employed in both the photoelectric artifact and optogenetic experiments to facilitate direct comparison and analysis of the results.

### Preparation of the brain slices for the illumination map characterization

The procedure to prepare the fixed brain slices presented below is approved by the Animal Experiments Inspectorate under the Danish Ministry of Food, Agriculture, and Fisheries, and all procedures adhere to the European guidelines for the care and use of laboratory animals, EU directive 2010/63/EU. A C57BL/6 mouse was anesthetized by the intraperitoneal injection of 0.01mL sodium pentobarbital (50 mg/kg). Once deep anesthesia was successfully achieved, the chest of the animal was opened in order to expose the heart. A needle, attached to a peristaltic pump (minipuls3, Gilson) via a catheter, was carefully inserted into the left ventricle of the heart before a small incision at the vena cava inferior was cut with fine surgery scissors to serve as the outlet of the blood during the perfusion. After 30 mL of 1X phosphate-buffered saline and 30 mL 4% paraformaldehyde (PFA) were perfused in succession, the animal was decapitated with a guillotine. The brain was dissected out and immersed in ice-cold 4% paraformaldehyde (PFA). The brain samples were incubated in PFA overnight and then with 30% sucrose solution for 2 days. The fixed brain was then cut into slices with a mouse brain matrice (coronal, 0.5mm cut, AgnTho’s). To prevent any movement of the brain during the cutting process, 3% (w/v) agarose gel was used to fill all the gaps between the stainless steel matrice and the brain tissue. Brain slices with a thickness of 500 µm were cut to measure the illumination map of the developed mAxialtrodes.

### Ethical approval and animal handling for *in vivo* experiments

Animal experiments were conducted according to the United Kingdom Animal (Scientific Procedures) Act 1986, and approved by the Animals in Scientific Research Unit at the Home Office. Animals were housed in a 12 h/12 h dark/light cycle, and food and water were given ad libitum. The animals were group-housed prior to headbar surgery, but after this, animals were individually housed.

### Virus injection

All following surgical procedures described were conducted aseptically. The C57BL/6 mice were anesthetized using isoflurane (988–3245, Henry Schein) followed by head shaving and skin disinfection. Then, the heads of the animals were fixed in a stereotaxic frame (David Kopf Instruments) by ear bars while a heating pad was used to maintain their body temperature. The body of the animal was covered with a sterile drape to minimize the temperature fluctuation during the surgery. Viscotears were applied (Bausch + Lomb). Buprenorphine (0.5 mg per/kg, Ceva) and Metacam (15 mg/kg, Boehringer) were subcutaneously injected for pain relief. After the animal lost its pedal reflex response, the skin on the top of the head was cut to expose the skull with a surgical scalpel, and a surgery drill was used to expose the cortex surface in the left hemisphere of the mouse brain for virus injection. 1uL of AAV9-pAAV-CaMKIIa-hChR2(H134R)-EYFP (Addgene) was injected into the visual cortex (coordinates: anteroposterior: -2.3 mm, mediolateral: 1.8 mm dorsoventral: 0.3 and 0.55 mm) using a stereotaxic frame at the speed of 75 nL/min.

### Surgeries for head bar attachment

Following injection of virus, a plate on the surface of the skull for the fixation of the animal’s brain with the head bar on the Neurostar system during subsequent awake in vivo experiments. After the skull was cleaned and dried, a small hole on the left hemisphere of the animal’s head was created by drilling (RA1 008, Henry Schein) for a metal support screw (00-96×3-32, Plastics One). The head plate was attached and strengthened using vetbond (1469SB, 3M), and dental cement, respectively flowed by the covering of the skull with silicone (Kwik-Cast, WPI). Saline was administered prior to recovery or every 45 min depending on the length of surgery.

Mice were checked daily to ensure recovery. After at least 5 days of recovery, habituation was performed by placing the mouse in the Neurotar frame for increasing periods of time (15–60 min) over several days.

### Craniotomy for the mAxialtrode implantation

A craniotomy was performed on the day of the in vivo experiment. After the anesthesia using isoflurane and the administration of pain medication and intramuscular dexamethasone (1 mg/kg, intramuscular; 7247104, MSD Animal Health), A large (2×2 mm^2^) craniotomy was performed above the somatosensory and visual cortex on the left hemisphere. Cold-buffered saline was continually applied to the craniotomies throughout the experiment. Then, the exposed dura was covered with buffered saline, a piece of sterilized Sylgard (∼200 μm thick), and a silicone layer. After two hours, the head of the animal was fixed to the Neurotar frame and the craniotomies were exposed by removal of the silicone and Sylguard layers. The tip of the mAxialtrode was lowered using a micromanipulator into the visual cortex or the hippocampus of the animal for neural stimulation or recordings. After the *in vivo* experiment (1-2hrs), sodium pentobarbital was administered intraperitoneally to humanely cull the animal prior to extraction of the brain for post-hoc analysis.

### Chronic biocompatibility evaluation of the mAxialtrodes

An immunohistochemistry experiment was carried out to evaluate the biocompatibility difference between the mAxialtrodes and conventional flat end-facet optical fibers. During probe implantation, a stereotaxic robot and drill were used to make two small holes in the skull in each hemisphere of mice (anterioposterior -2 mm, mediolateral 1.5 mm, dorsoventral 2 mm from dura mater). The mAxialtrodes were implanted in the left hemisphere while the fiber with the same diameter but a flat end-facet was implanted in the right hemisphere. The skull was then sealed by cyanoacrylate adhesive and dental cement. After the surgery, the mice were orally treated with buprenorphine (0.2 mg mixed with 1 g Nutella) every 12 hours for three days. Furthermore, Carprofen (5 mg/kg) was administered subcutaneously every 12 hours for 5 days. The conditions of the animals were monitored 2–3 times per day over the first 3 days and once daily for the following week.

Four weeks after the fiber implantation, the mice were perfused and their brains were removed, and cryoprotected with the same procedure as described above when getting brain slices for the illumination map measurement. A cryostat (CM3050S, Leica) was used to slice the brain with a thickness of 50 µm. After being washed in PBS and preincubated in 0.3% Triton X-100 (PBST) for 15 mins, the slices were then incubated in a blocking solution consisting of 3% bovine serum albumin (BSA) for 1 h at 25 °C. Primary antibodies (1:1000 dilution, rabbit polyclonal, Wako 019-19741) were applied to the slice for overnight incubation at 4 °C. Afterwards, the slices were washed three times with 0.1 % PBST. Then, secondary antibodies (goat anti-rabbit, Alexa Fluor 488,Invitrogen) together with 4’,6-Diamidino-2-Phenylindole, Dihydrochloride (DAPI, Invitrogen, D1306) was applied at a concentration of 1:1000. After the incubation in dark room for 1h at 25 °C, the slices were rewashed three times with 0.1 % PBST. Finally, the glass slides were mounted using DAKO mounting medium and imaged using a Zeiss Axioscan Z1 microscope (×10 magnification).

Images were loaded at full resolution in Zeiss ZEN software and two different square ROIs with a dimension of 500 × 500 µm centered around the tip of the mAxialtrodes and the flat end-facet fibers were cropped. After discarding the images found to be out of focus or presenting too large artifacts, a threshold was applied to eliminate the signal from out-of-plane cells, and the maximum pixel value was adjusted to increase contrast. The analysis of the images was achieved by the Fiji open-source image processing package, based on ImageJ2. After the marker and DAPI channels were separated, the mean fluorescence intensity of the Iba-1 marker was calculated.

## Data processing

The data processing and analysis were performed in the Matlab 2022b suite. The SNR was calculated by dividing the median amplitude of all spike events surpassing the threshold by the threshold value itself ^37^. Here the threshold for spike detection was set as ^55^

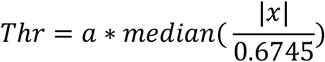

where |*χ*| represents the amplitude of the high-passed data, and *a* represents a factor for different threshold levels. This factor was set to a = 3 for the standard procedure and was varied between 2 and 4 for an iterative procedure. We use a = 3 in this work.

In the PCA analysis, the principal component coefficients are obtained by the singular value decomposition (SVD) algorithm. The clusters were sorted based on fitting the Gaussian mixture model to the data. The Gaussian mixture model was initialized with the k-means++ algorithm and used diagonal and identical covariance matrices to fit the data.

## Acknowledgments

C.M. acknowledge funding from Lundbeck Foundation Multi-BRAIN project R276-2018-869, EIC Pathfinder project Move2Treat (GAP 101130161) and R380-2021-1171, M.M. acknowledge funding from Villum Fonden (36063)

## Author contributions

K.S. and C.M. conceptualized the microfluidic axialtrode and designed the experiments. K.S. developed the microfluidic axialtrode and characterized its optical and electrical properties. K.S. and A.B. conducted the drug delivery characterization. M.M. and A.B. designed and fabricated the biocompatible scaffold. K.S. and G.L. performed brain slicing and illumination mapping. K.S. and G.L. conducted the immunohistochemistry experiments. N.C. and D.R. performed mouse surgeries for in vivo experiments. K.S., N.C., and D.R. conducted EEG recordings, and K.S. and M.M. analyzed the EEG data. K.S. drafted the manuscript and prepared the figures. All authors contributed to manuscript editing. C.M., R.B., and R.W. supervised the project. C.M., M.M., R.B., and R.W. secured funding.

### Data and materials availability

All data needed to evaluate the conclusions in the paper are present in the paper and/or the Supplementary Materials. Additional data related to this paper may be requested from the authors.

## References

1. Ramezani, M., Ren, Y., Cubukcu, E. & Kuzum, D. Innovating beyond electrophysiology through multimodal neural interfaces. Nat Rev Electr Eng 2, 42–57 (2025).

2. Rivnay, J., Wang, H., Fenno, L., Deisseroth, K. & Malliaras, G. G. Next-generation probes, particles, and proteins for neural interfacing. Science Advances 3, e1601649 (2017).

3. Delcasso, S., Denagamage, S., Britton, Z. & Graybiel, A. M. HOPE: Hybrid-Drive Combining Optogenetics, Pharmacology and Electrophysiology. Front. Neural Circuits 12, (2018).

4. Sharma, K. et al.. Multifunctional optrode for opsin delivery, optical stimulation, and electrophysiological recordings in freely moving rats. J. Neural Eng. 18, 066013 (2021).

5. Chapman, C. A. R., Goshi, N. & Seker, E. Multifunctional Neural Interfaces for Closed-Loop Control of Neural Activity. Advanced Functional Materials 28, 1703523 (2018).

6. Shin, H. et al.. Neural probes with multi-drug delivery capability. Lab on a Chip 15, 3730–3737 (2015).

7. Zheng, N. et al.. Multifunctional Fiber-Based Optoacoustic Emitter as a Bidirectional Brain Interface. Advanced Healthcare Materials n/a, 2300430.

8. Zhang, Y. et al. Submillimeter Multifunctional Ferromagnetic Fiber Robots for Navigation, Sensing, and Modulation. Advanced Healthcare Materials n/a, 2300964.

9. Chin, A. L. et al.. Implantable optical fibers for immunotherapeutics delivery and tumor impedance measurement. Nat Commun 12, 5138 (2021).

10. Jiang, S. et al.. Spatially expandable fiber-based probes as a multifunctional deep brain interface. Nat Commun 11, 6115 (2020).

11. Anikeeva, P. et al.. Optetrode: a multichannel readout for optogenetic control in freely moving mice. Nat Neurosci 15, 163–170 (2012).

12. Park, S. et al.. One-step optogenetics with multifunctional flexible polymer fibers. Nature Neuroscience 20, 612–619 (2017).

13. Park, S., Loke, G., Fink, Y. & Anikeeva, P. Flexible fiber-based optoelectronics for neural interfaces. Chemical Society Reviews 48, 1826–1852 (2019).

14. Xu, P., Chen, A., Li, Y., Xing, X. & Lu, H. Medial prefrontal cortex in neurological diseases. Physiological Genomics 51, 432–442 (2019).

15. Bonaccini Calia, A. et al. Full-bandwidth electrophysiology of seizures and epileptiform activity enabled by flexible graphene microtransistor depth neural probes. Nat. Nanotechnol. 17, 301–309 (2022).

16. Sileo, L. et al.. Tapered Fibers Combined With a Multi-Electrode Array for Optogenetics in Mouse Medial Prefrontal Cortex. Frontiers in Neuroscience 12, (2018).

17. Chen, M. et al.. Self-powered multifunctional sensing based on super-elastic fibers by soluble-core thermal drawing. Nature Communications 12, 1416 (2021).

18. Cariou, J. M., Dugas, J., Martin, L. & Michel, P. Refractive-index variations with temperature of PMMA and polycarbonate. Appl. Opt., AO 25, 334–336 (1986).

19. Keck, C. H. C., Rommelfanger, N. J., Ou, Z. & Hong, G. Bioinspired nanoantennas for opsin sensitization in optogenetic applications: a theoretical investigation. Multifunct. Mater. 4, 024002 (2021).

20. Ambrosi, C. M. & Entcheva, E. Optogenetic Control of Cardiomyocytes via Viral Delivery. in Cardiac Tissue Engineering: Methods and Protocols (eds. Radisic, M. & Black III, L. D.) 215–228 (Springer, New York, NY, 2014). doi:10.1007/978-1-4939-1047-2_19.

21. Chen, S. Optical modulation goes deep in the brain. Science 365, 456–457 (2019).

22. Zhou, Z. et al.. Towards high-power mid-IR light source tunable from 3.8 to 4.5 µm by HBr-filled hollow-core silica fibres. Light Sci Appl 11, 15 (2022).

23. Stefani, A., Yuan, W., Markos, C. & Bang, O. Narrow Bandwidth 850-nm Fiber Bragg Gratings in Few-Mode Polymer Optical Fibers. IEEE Photonics Technology Letters 23, 660–662 (2011).

24. Fasano, A. et al.. Fabrication and characterization of polycarbonate microstructured polymer optical fibers for high-temperature-resistant fiber Bragg grating strain sensors. Opt. Mater. Express, OME 6, 649–659 (2016).

25. Tanaka, A., Sawada, H., Takoshima, T. & Wakaisuki, N. New Plastic Optical Fiber With Polycarbonate Core And Fluorescence-Doped Fiber For High Temperature Use. in Fiber Optic Systems for Mobile Platforms vol. 0840 19–28 (pnSPIE, 1987).

26. Yamashita, T. Y. T. & Kamada, K. K. K. Intrinsic Transmission Loss of Polycarbonate Core Optical Fiber. Jpn. J. Appl. Phys. 32, 2681 (1993).

27. Park, S. et al.. Adaptive and multifunctional hydrogel hybrid probes for long-term sensing and modulation of neural activity. Nat Commun 12, 3435 (2021).

28. Dijk, G., Rutz, A. L. & Malliaras, G. G. Stability of PEDOT:PSS-Coated Gold Electrodes in Cell Culture Conditions. Advanced Materials Technologies 5, 1900662 (2020).

29. Yu, L. et al.. Optogenetic Modulation of Cardiac Sympathetic Nerve Activity to Prevent Ventricular Arrhythmias. Journal of the American College of Cardiology 70, 2778–2790 (2017).

30. Azadi, R. et al.. Image-dependence of the detectability of optogenetic stimulation in macaque inferotemporal cortex. Current Biology 33, 581–588.e4 (2023).

31. Messaritaki, E., Rudrapatna, S. U., Parker, G. D., Gray, W. P. & Jones, D. K. Improving the Predictions of Computational Models of Convection-Enhanced Drug Delivery by Accounting for Diffusion Non-gaussianity. Front. Neurol. 9, (2018).

32. Brusso, B. C. A Brief History of the Energy Conversion of Light [History]. IEEE Industry Applications Magazine 25, 8–13 (2019).

33. Buzsáki, G. et al.. Tools for Probing Local Circuits: High-Density Silicon Probes Combined with Optogenetics. Neuron 86, 92–105 (2015).

34. Yazdan-Shahmorad, A., Silversmith, D. B., Kharazia, V. & Sabes, P. N. Targeted cortical reorganization using optogenetics in non-human primates. eLife 7, e31034 (2018).

35. Mikulovic, S. et al.. On the photovoltaic effect in local field potential recordings. NPh 3, 015002 (2016).

36. Kozai, T. D. Y. & Vazquez, A. L. Photoelectric artefact from optogenetics and imaging on microelectrodes and bioelectronics: new challenges and opportunities. J. Mater. Chem. B 3, 4965–4978 (2015).

37. Drebitz, E., Schledde, B., Kreiter, A. K. & Wegener, D. Optimizing the Yield of Multi-Unit Activity by Including the Entire Spiking Activity. Front. Neurosci. 13, (2019).

38. MacKay, W. A. Synchronized neuronal oscillations and their role in motor processes. Trends in Cognitive Sciences 1, 176–183 (1997).

39. Puentes-Mestril, C., Roach, J., Niethard, N., Zochowski, M. & Aton, S. J. How rhythms of the sleeping brain tune memory and synaptic plasticity. Sleep 42, zsz095 (2019).

40. Lindén, H., Petersen, P. C., Vestergaard, M. & Berg, R. W. Movement is governed by rotational neural dynamics in spinal motor networks. Nature 610, 526–531 (2022).

41. Buzsáki, G. Theta Oscillations in the Hippocampus. Neuron 33, 325–340 (2002).

42. Sih, G. C. & Tang, K. K. Sustainable time and stability of hippocampal and cortical EEG theta waves. Theoretical and Applied Fracture Mechanics 62, 1–14 (2012).

43. Deisseroth, K. Optogenetics. Nat Methods 8, 26–29 (2011).

44. Bamann, C., Kirsch, T., Nagel, G. & Bamberg, E. Spectral Characteristics of the Photocycle of Channelrhodopsin-2 and Its Implication for Channel Function. Journal of Molecular Biology 375, 686–694 (2008).

45. Erofeev, A. et al.. Light Stimulation Parameters Determine Neuron Dynamic Characteristics. Applied Sciences 9, 3673 (2019).

46. Stiller, A. M. et al.. Mechanical considerations for design and implementation of peripheral intraneural devices. J. Neural Eng. 16, 064001 (2019).

47. Sui, K., Meneghetti, M., Berg, R. W. & Markos, C. Optoelectronic and mechanical properties of microstructured polymer optical fiber neural probes. Optics Express 31, 21563–21575 (2023).

48. Sharp, A. A., Ortega, A. M., Restrepo, D., Curran-Everett, D. & Gall, K. In Vivo Penetration Mechanics and Mechanical Properties of Mouse Brain Tissue at Micrometer Scales. IEEE Transactions on Biomedical Engineering 56, 45–53 (2009).

49. Kim, J. et al.. T-DOpE probes reveal sensitivity of hippocampal oscillations to cannabinoids in behaving mice. Nat Commun 15, 1686 (2024).

50. Canales, A. et al.. Multifunctional fibers for simultaneous optical, electrical and chemical interrogation of neural circuits in vivo. Nature Biotechnology 33, 277–284 (2015).

51. Lee, L.-H. & Chen, W.-C. High-Refractive-Index Thin Films Prepared from Trialkoxysilane-Capped Poly(methyl methacrylate)-Titania Materials. Chem. Mater. 13, 1137–1142 (2001).

52. Trujillo-Pisanty, I., Sanio, C., Chaudhri, N. & Shizgal, P. Robust optical fiber patch-cords for in vivo optogenetic experiments in rats. MethodsX 2, 263–271 (2015).

53. Bouchekioua, Y. et al.. Striatonigral direct pathway activation is sufficient to induce repetitive behaviors. Neuroscience Research 132, 53–57 (2018).

54. Maglie, E. et al.. Ray tracing models for estimating light collection properties of microstructured tapered optical fibers for optical neural interfaces. Opt. Lett., OL 45, 3856–3859 (2020).

55. Quiroga, R. Q., Nadasdy, Z. & Ben-Shaul, Y. Unsupervised Spike Detection and Sorting with Wavelets and Superparamagnetic Clustering. Neural Computation 16, 1661–1687 (2004).

