## Supplementary material for "Multimodal layer-crossing interrogation of brain circuits enabled by microfluidic axialtrodes": Figures and tables

1. **Supplementary Figures**


| | |
|---|---|
| **Supplementary Figure 1** | The drawn PC/PMMA polymer optical fibers |
| **Supplementary Figure 2** | Mechanical cutting method for introducing an angle at the tip of optical fibers |
| **Supplementary Figure 3** | Different resolution targets of Formlabs Biomed Clear |
| **Supplementary Figure 4** | The 3D printed scaffolds for the mAxialtrode |
| **Supplementary Figure 5** | Two mice with the implanted mAxialtrode devices (design 2) on their heads |
| **Supplementary Figure 6** | The model in ray-tracing software Zemax OpticStudio for measuring the illumination map of the optical fibers in the medium of the brain |
| **Supplementary Figure 7** | The 3D models and their side views in ray-tracing software Zemax OpticStudio to illustrate the light emission angle variation with the decreasing of the optical fiber tip angle from 90° to 15° |
| **Supplementary Figure 8** | The illumination map of the angled tip fiber (30°) in brain slices |
| **Supplementary Figure 9** | Impedance spectroscopy of the eight integrated tungsten electrodes in the mAxialtrode |
| **Supplementary Figure 10** | The drug delivery characterization of the mAxialtrode |
| **Supplementary Figure 11** | Statistical significance analysis of the IHC results at the implantation sites |
| **Supplementary Figure 12** | The schematic of the mAxialtrode device connection |
| **Supplementary Figure 13** | The full photoelectric artifacts recordings during pulse light stimulation in PBS |
| **Supplementary Figure 14** | The power spectrum of the light-evoked artifacts collected by the distributed eight electrodes in the mAxialtrode |
| **Supplementary Figure 15** | The data analysis with highpass filter to minimize the photoelectric artifacts |
| **Supplementary Figure 16** | Full electrophysiology recordings by the mAxialtrode devices |
| **Supplementary Figure 17** | The theta rhythm component of the electrophysiology recordings by the mAxialtrode device |

| | |
|---|---|
| **Supplementary Figure 18** | Typical electrophysiological recordings using the mAxialtrode device in optogenetic experiments demonstrate brain extracellular responses to light-induced activity |
| **Supplementary Figure 19** | Typical 1s electrophysiological recordings in optogenetic experiments to show the details of the eight channels' traces in Supplementary Fig. 18 |
| **Supplementary Figure 20** | Optogenetic control of action potential firing in the mouse brain |
| **Supplementary Figure 21** | The evoked neural activity amplitude from all eight electrodes during optical stimulation with different pulse widths (10 ms, 20 ms, and 50 ms). |

### 2. Supplementary Tables

| | |
|---|---|
| **Supplementary Table 1** | Print settings for Formlabs Biomed Clear resin on Phrozen Mighty 8K. |
| **Supplementary Table 2** | The SNR of the spikes under different light pulse stimulation |

### 3. Supplementary Videos

| | |
|---|---|
| **Supplementary Video 1** | The mice with the implanted mAxialtrode device |
| **Supplementary Video 2** | The schematic of the neural activity recording process by the mAxialtrode |

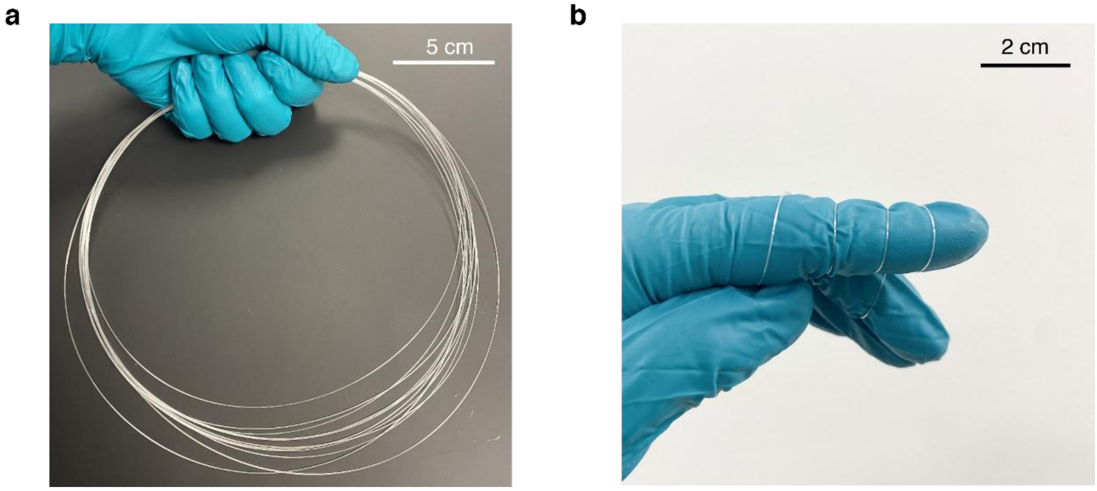


**Supplementary Figure 1. The drawn PC/PMMA polymer optical fibers**. **a,** Tens of meters of polymer optical fiber were produced with the thermal drawing method. **b,** The fibers have high flexibility, allowing them to be wrapped around a finger without breaking.

a b

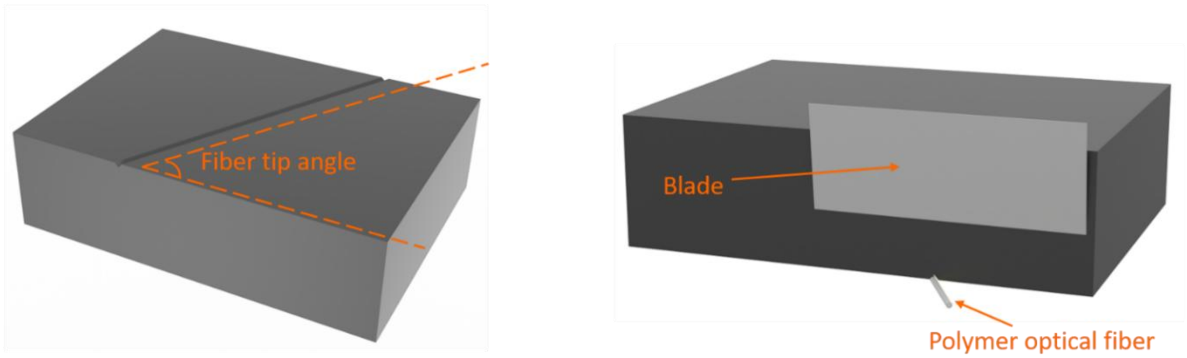


**Supplementary Figure 2. Mechanical cutting method for introducing an angle at the tip of optical fibers. a,** The groove was machined at one surface of a stainless steel block. The angle between the groove and the edge of the block was chosen based on the target fiber tip angle. **b,** The polymer optical fiber was placed in the groove, and the cutting was achieved by sliding the blade down along the side face of the stainless steel block.

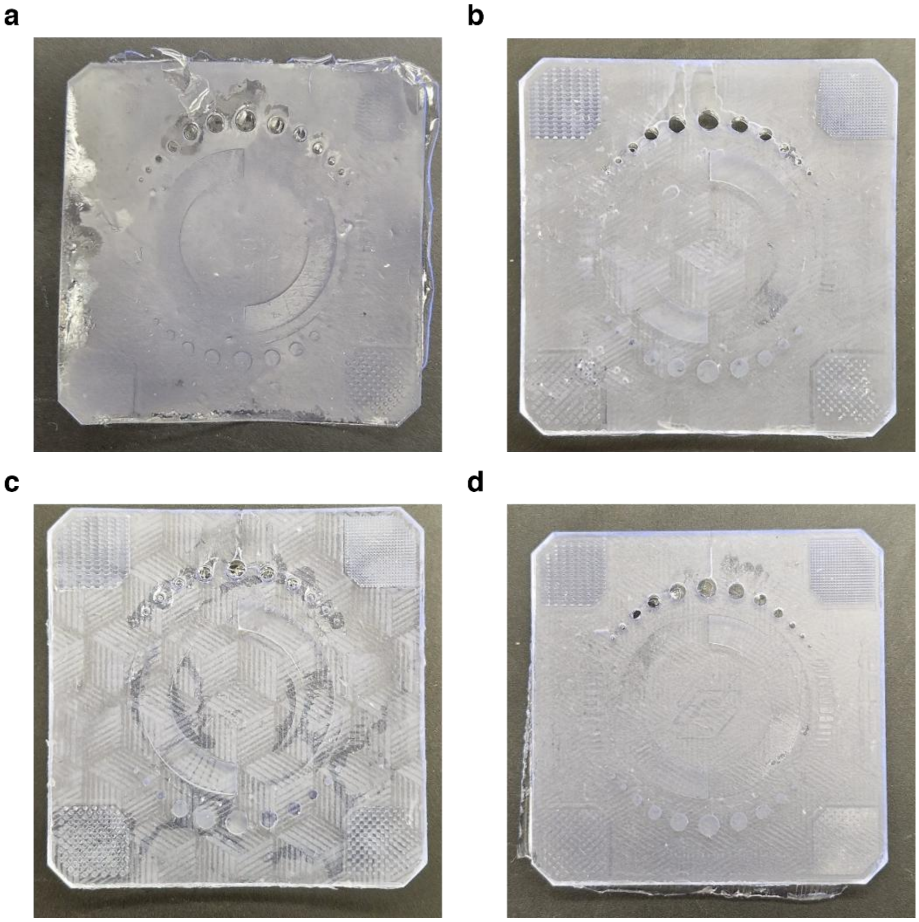

**Supplementary Figure 3. Different resolution targets of Formlabs Biomed Clear: a,** Very under-exposed resin, a strong lack of detail, as the resin did not solidify. **b,** Insufficient exposure leading to the exposure target not sticking completely and resin leaking under the bottom. **c,** Overexposure at the bottom, leading to too much resin hardening at the bottom of the exposure target. **d,** A successful print is slightly overexposed at the bottom.

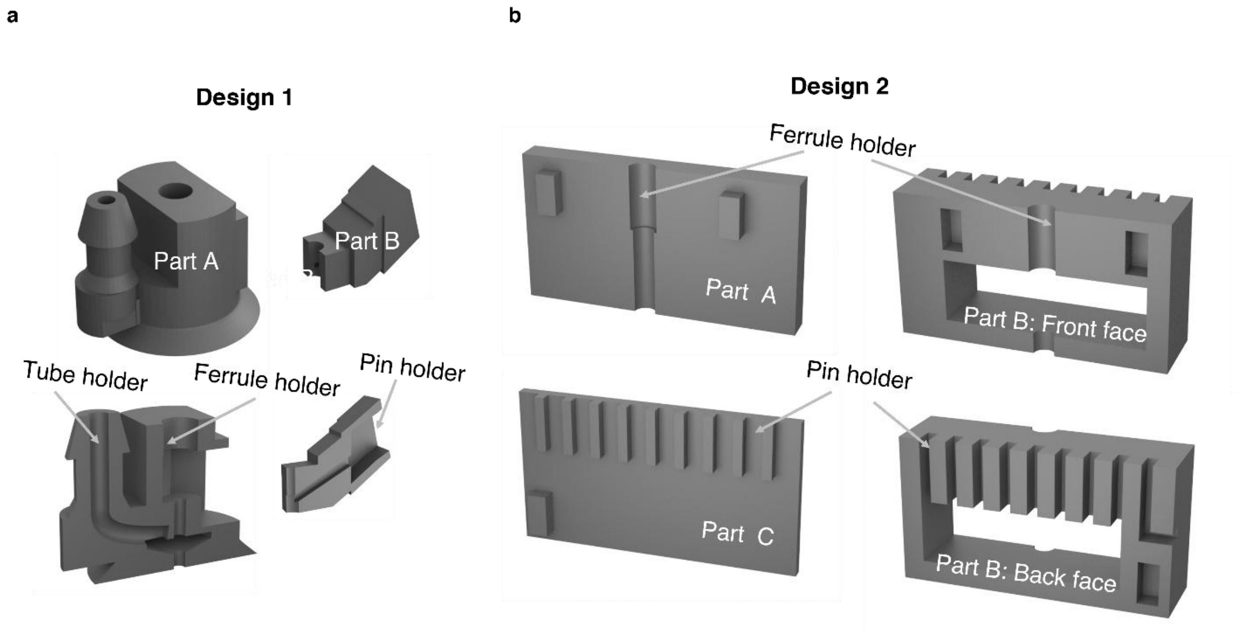


**Supplementary Figure 4. The 3D printed scaffolds for the mAxialtrode. a,** The detailed structures of the design 1 scaffold. Part A is the main body of the scaffold, which includes a tube holder and an optical ferrule holder. When electrodes are integrated into the mAxialtrodes, the metal wires can be connected with electric pins by Part B. **b,** The detailed structures of the three parts of the mAxialtrode scaffold in design 2. The assembling of parts A and B can fix the position of the ferrule on the mAxialtrode, while the assembling of parts B and C can fix the position of the eight tungsten wires and the grounding wire to a 1.27 mm pin header (9 pins in total).In the assembly process, firstly, the position of the ferrule attached mAxialtrode was fixed by assembling part A to the front face of part B. Secondly, a tungsten wire, which would work as the grounding wire in the in vivo test, was placed into the rightmost slot in the back face of Part B, followed by the placement of the eight tungsten wires in the mAxialtrode to the rest eight slots in the back face of Part B. Then, a 9-pin header was inserted into the 9 slots in the back face of Part B. Finally, Part C was assembled to the back face of Part B. In this design, instead of connecting the metal wires to the pins by soldering, we achieved a robust electrode-pin connectorization through the interference fit between the pin holders in Part B and Part C, ensuring a secure connection between the tungsten wires and the pins in the nine slots.

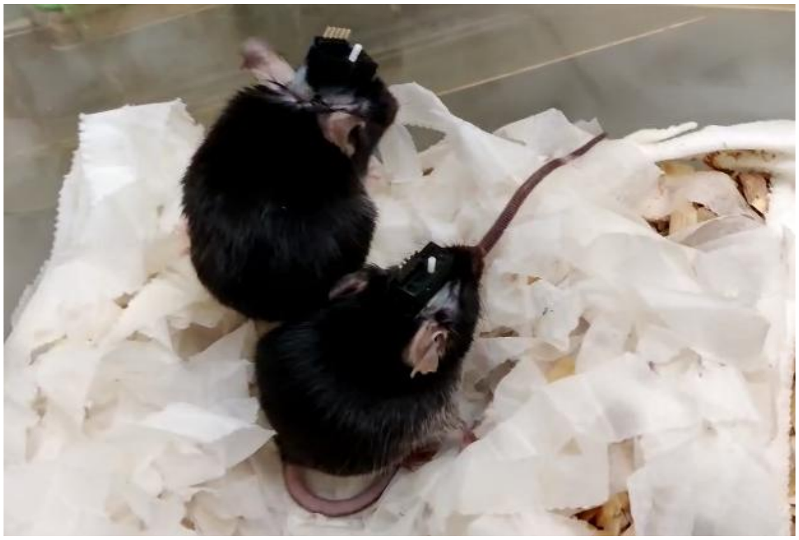

**Supplementary Figure 5. Two mice with the implanted mAxialtrode device (design 2) on their heads.**

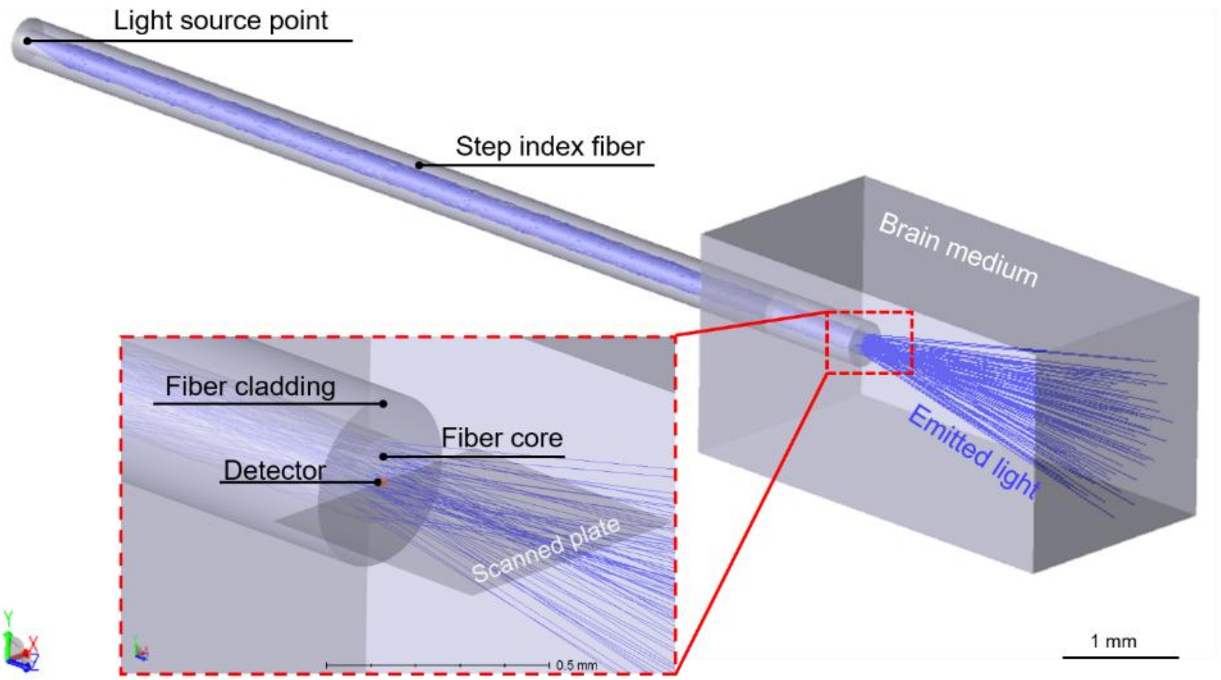


**Supplementary Figure 6. The model in ray-tracing software Zemax OpticStudio for measuring the illumination map of the optical fibers in the medium of the brain.**

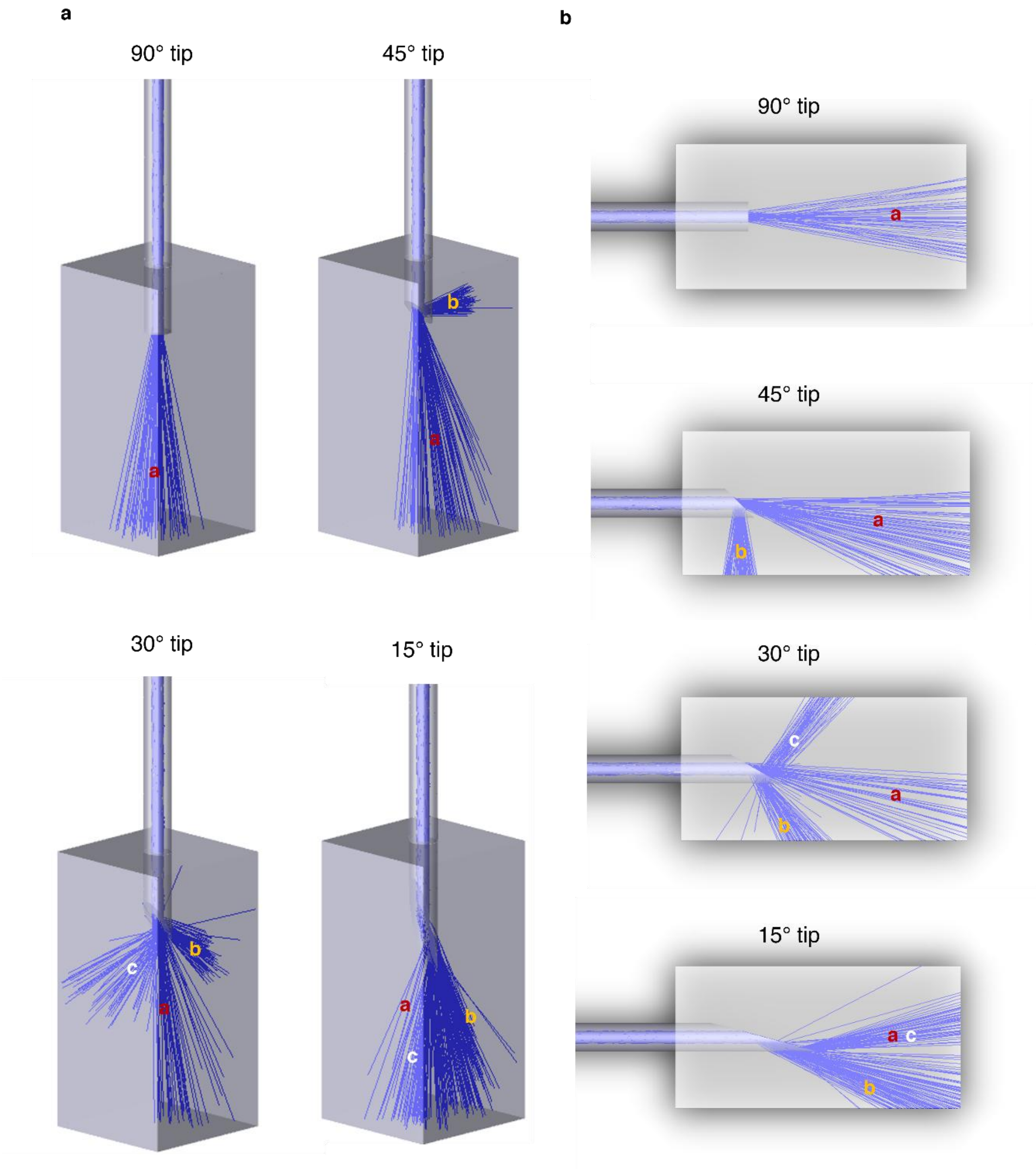


**Supplementary Figure 7. The 3D models and their side views in ray-tracing software Zemax OpticStudio to illustrate the light emission angle variation with the decreasing of the optical fiber tip angle from 90° to 15°.** When the tip of the optical fiber is flat (90°), all the rays are emitted to the direction along the length of the fiber (*cluster a*). When a smaller angle is introduced to the tip of the fiber (45°), some of the rays emitted from the fiber end facet are reflected by the interface between the fiber core and brain medium and propagated to the side of the fiber (*cluster b*). As the tip of the fiber becomes shaper (30°), some of the rays in cluster b are reflected by the interface between fiber cladding and the brain medium and propagated to the other side of the fiber (*cluster c)*. The angle between *cluster b* and *cluster c* can be reduced with the further decrease of the fiber tip (15°).

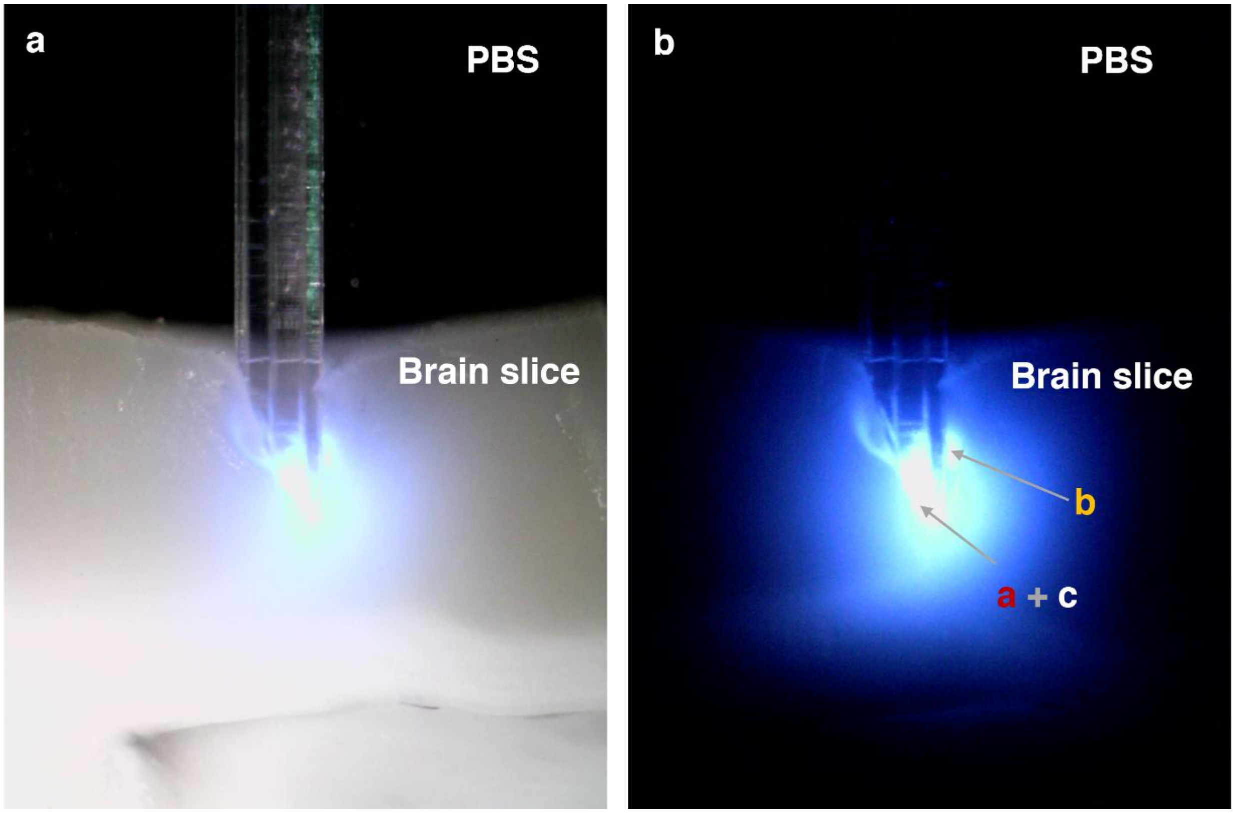


**Supplementary Figure 8. The illumination map of the angled tip fiber (30°) in brain slices. a,** The tip of the fiber was inserted into the brain slices under microscope light for position conformation. **b,** without microscope light. It can be seen that most of the blue light was emitted from the tip of the fiber, which can match the simulation result from Zemax OpticStudio. However, the illumination map behaved in a strong inhomogeneous because of the randomly squeezed brain tissue during the mAxialtrode insertion. To reduce these squeezed-tissue-artifacts in the illumination map, the brain slice was cut in the rostral-caudal direction, and the flat upper surface of the cortex was positioned in touch with the end facet of the angled tip fiber as seen in Fig. 2b.

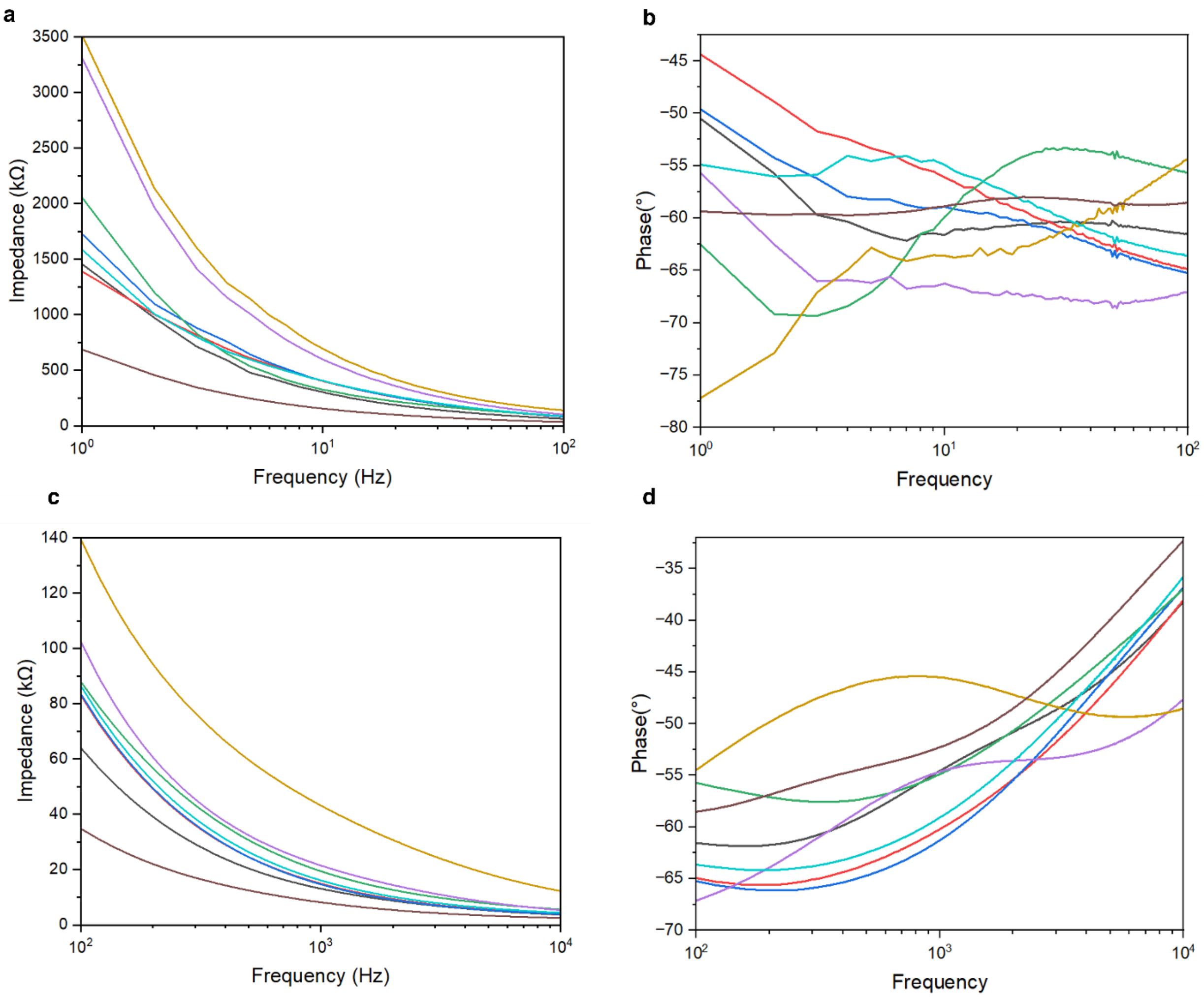


**Supplementary Figure 9. Impedance spectroscopy of the eight integrated tungsten electrodes in the mAxialtrode.** To enhance visualization, the broadband results are divided into two sub-bands: 1-100 Hz (**a** and **b**) and 100-10 kHz (**c** and **d**). **a,** The measured impedance for each of the eight electrodes in design 2 within the frequency band of 1-100 Hz, different colors are used to represent the measured results from each of the eight electrodes. **b,** The phase measured for each of the eight electrodes in design 2 within the 1-100 Hz frequency band**. c,** The measured impedance for each of the eight electrodes in design 2 within the frequency band of 100-10 kHz. **d,** The phase measured for each of the eight electrodes in design 2 within the 100-10 kHz frequency band**.** The measured impedance of all of eight electrodes in the mAxialtrode is notably below the threshold for neural activity detection (1 MΩ at 1 kHz).

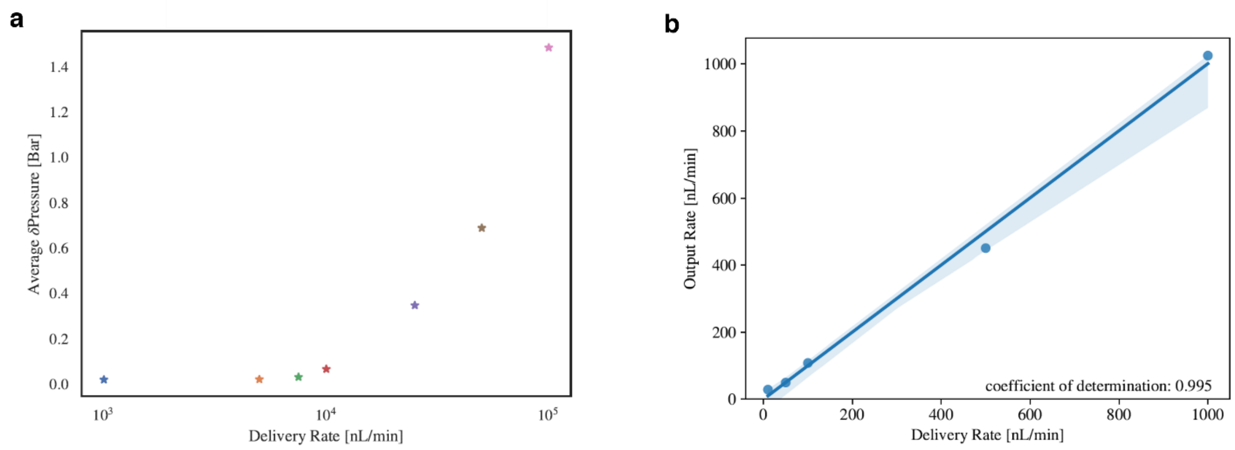


**Supplementary Figure 10. The drug delivery characterization of the mAxialtrode. a,** The average pressure of fiber in the microfluidic channels at different delivery rates from 1000 nL/min to 100,000 nL/min within 20 mins. **b,** the output rate compared to the delivery rate, showing confidence of determination as 0.995.

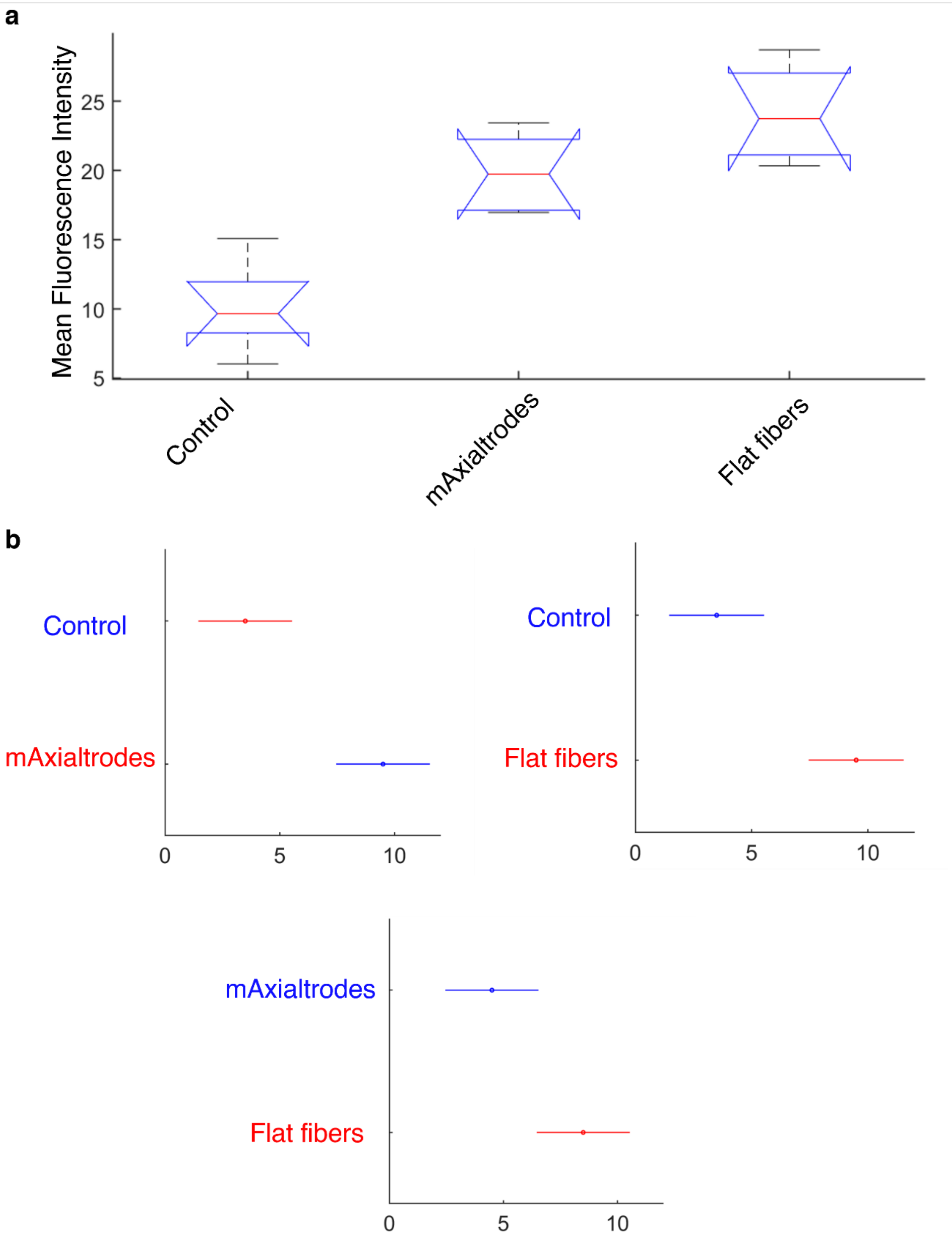


**Supplementary Figure 11. Statistical significance analysis of the IHC results at the implantation sites. a,** The Kruskal-Wallis test suggests statistically significant ($p<0.05$) differences between the control, mAxialtrode, and flat fiber groups. **b,** Pairwise Tukey honest significance tests have been performed for protection against type I errors. The null hypothesis can be rejected at $p<0.05$ by the tests of the Control-flat fiber and control-mAxialtrodes pairs. By contrast, the mAxialtrodes-flat fiber pairs test results in a slightly higher p-value (0.0547).

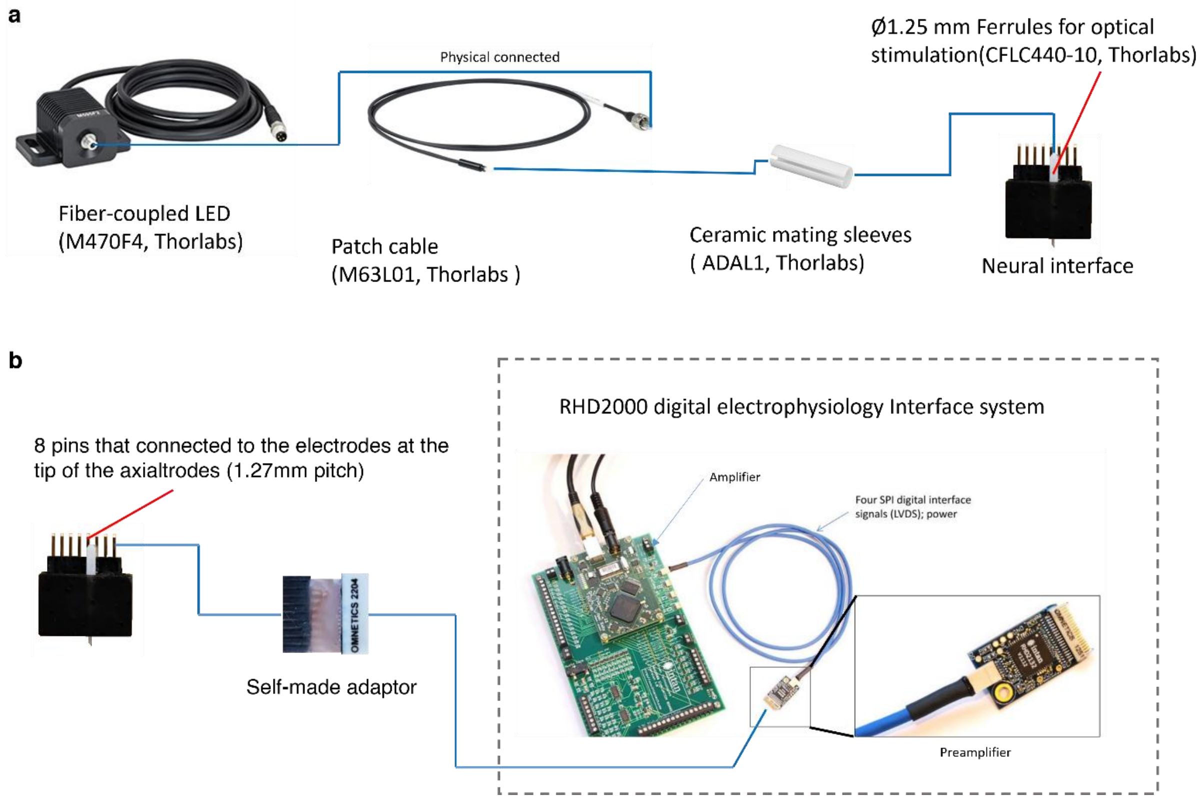


**Supplementary Figure 12. The schematic of the mAxialtrode device connection. a,** optical stimulation and **b,** electrophysiology recordings (The physical connections between the components' ports are represented by blue lines).

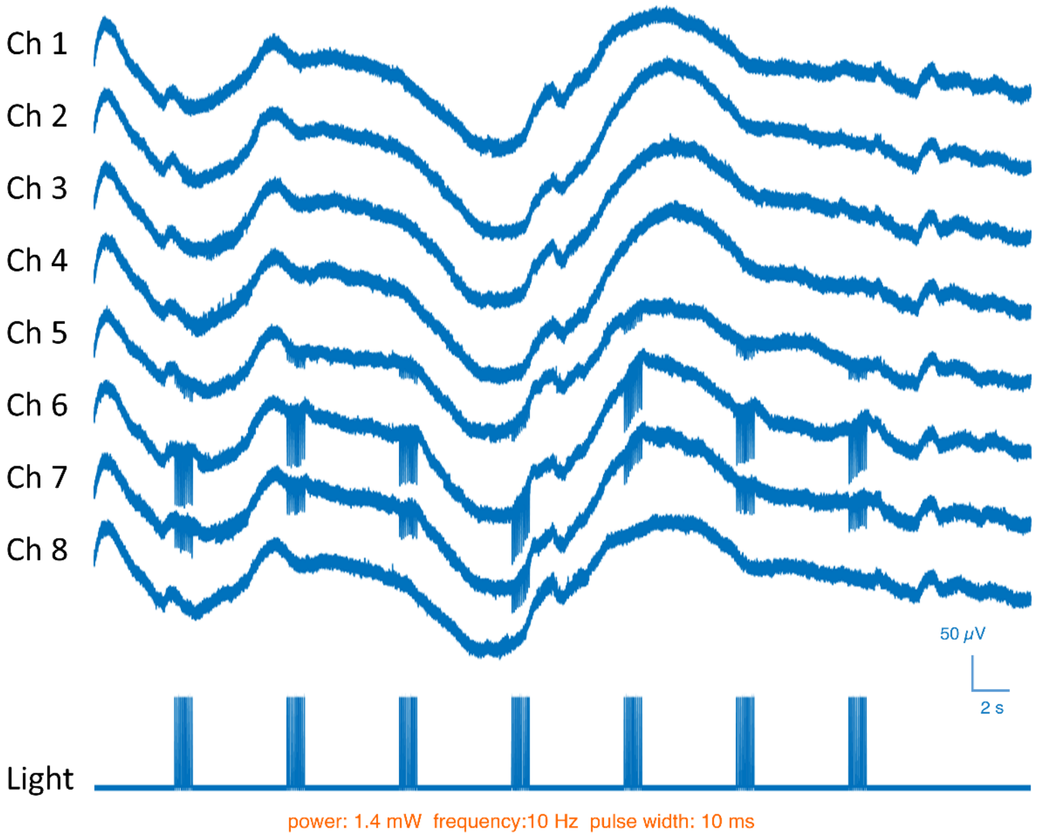


**Supplementary Figure 13. The full photoelectric artifacts recordings during pulse light stimulation in PBS.** The three electrodes near the tip of the mAxialtrode (Ch5, Ch6, and Ch7) behave with light-induced electric signals, while the light stimulation has a negligible effect on the other electrodes.

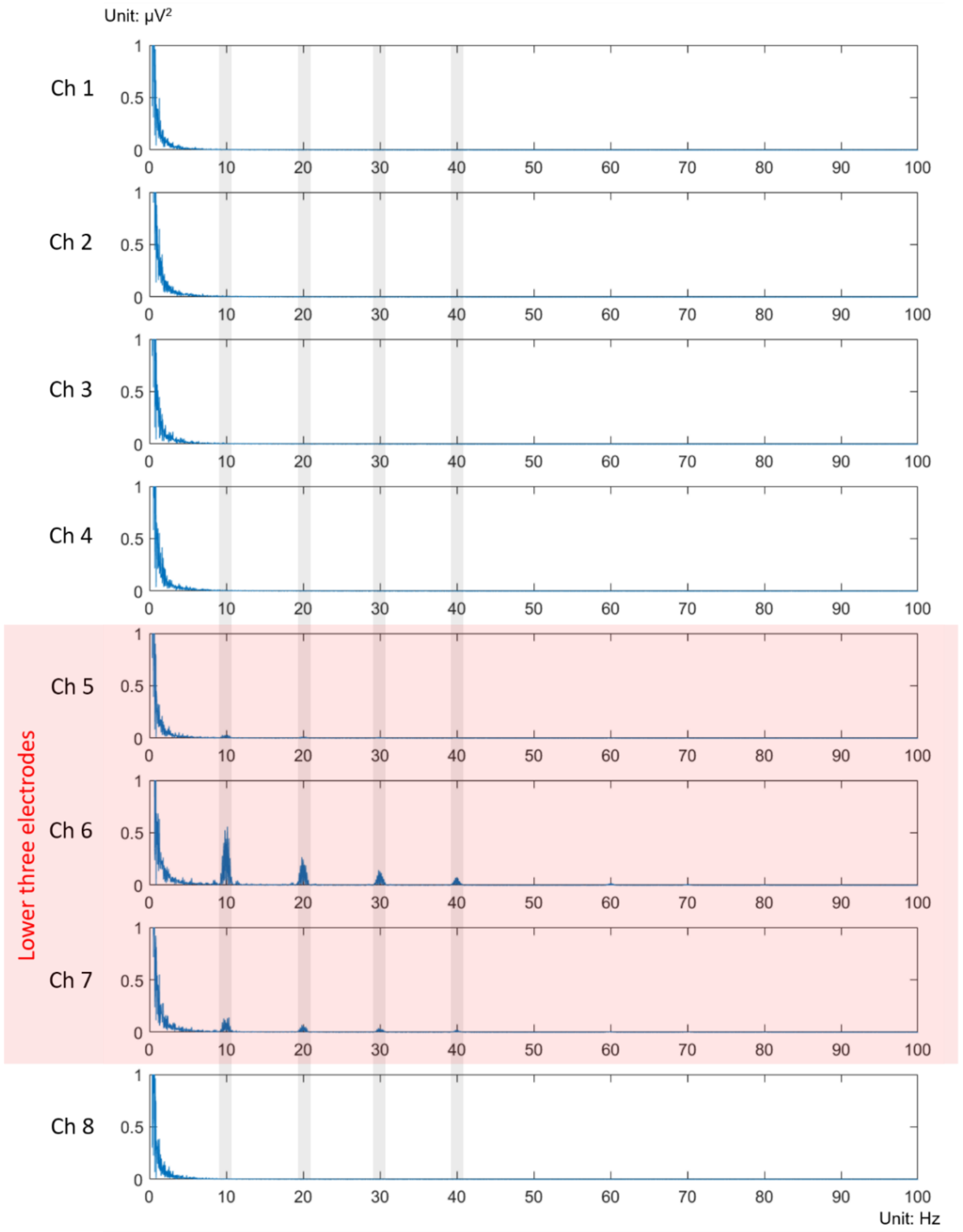


**Supplementary Figure 14. The power spectrum of the light-evoked artifacts collected by the distributed eight electrodes in the mAxialtrode.** It is clear that the photoelectric artifacts have the same frequency as the stimulation light (10 Hz) and its overtones (20, 30, and 40 Hz). The power of the photoelectric artifacts keeps decreasing at a higher frequency band. This makes it possible to minimize the photoelectric artifacts by using a high-pass filter. In addition, the light-evoked artifacts are mainly observed in the lower three electrodes in the neural interface. This is due to that most of the light propagating in the fiber core would be emitted from the tip of the fiber when a small tip angle is introduced to the mAxialtrode, as illustrated in fig. S6. Therefore, only the lower three electrodes have been illuminated by the stimulation light in the PBS medium.

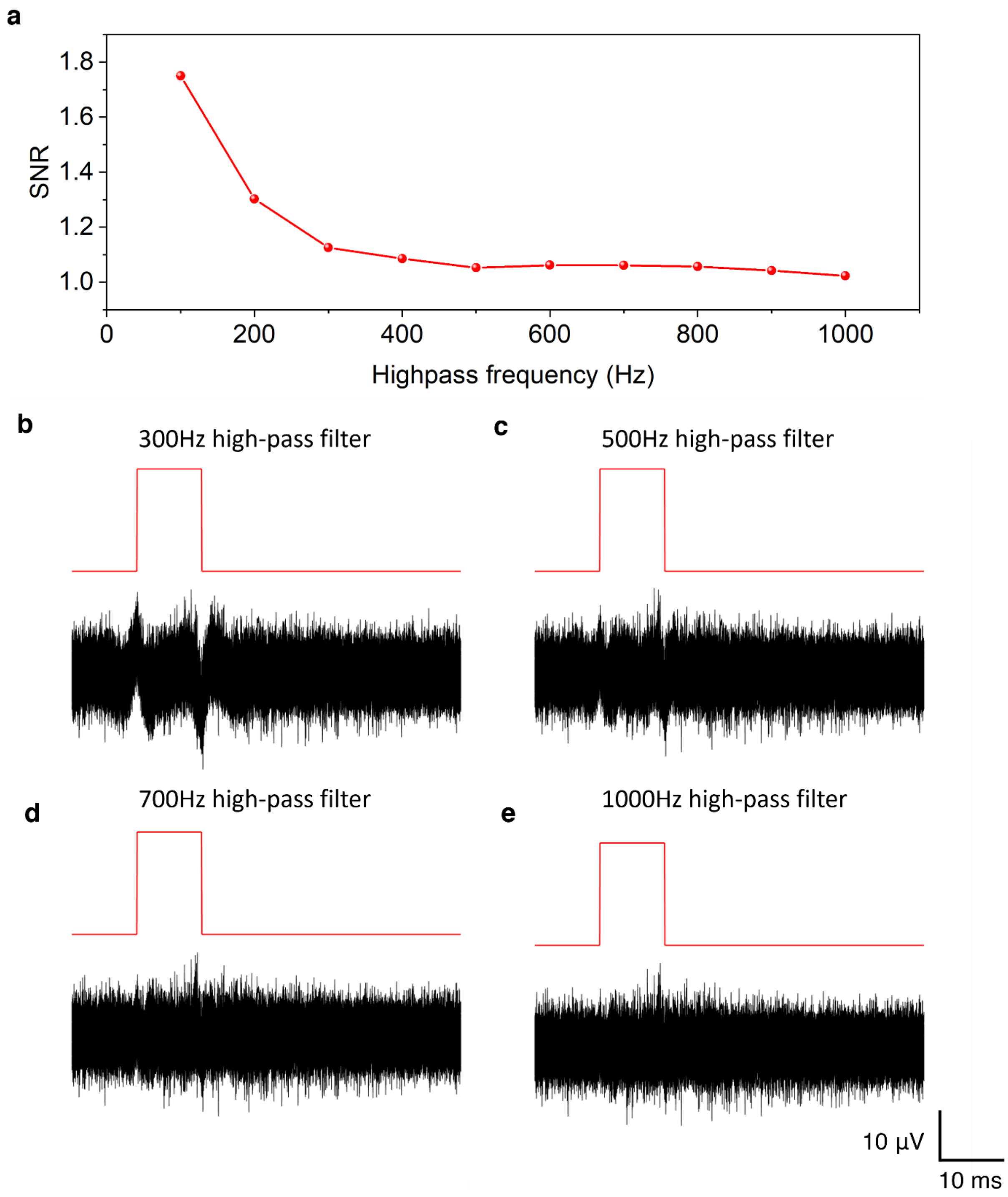


**Supplementary Figure 15. The data analysis with highpass filter to minimize the photoelectric artifacts. a,** Different highpass filters are applied to the recorded data from Ch6 (which exhibits the strongest photoelectric artifacts) in Supplementary Fig. 13 to calculate the photoelectric artifacts SNR under light stimulation. It can be seen that there are some residential photoelectric artifacts after using a 300 Hz highpass filter in **b**, while the artifacts can be minimized when increasing the frequency of the highpass filter to 500 Hz **c**, **d**, and **e**.

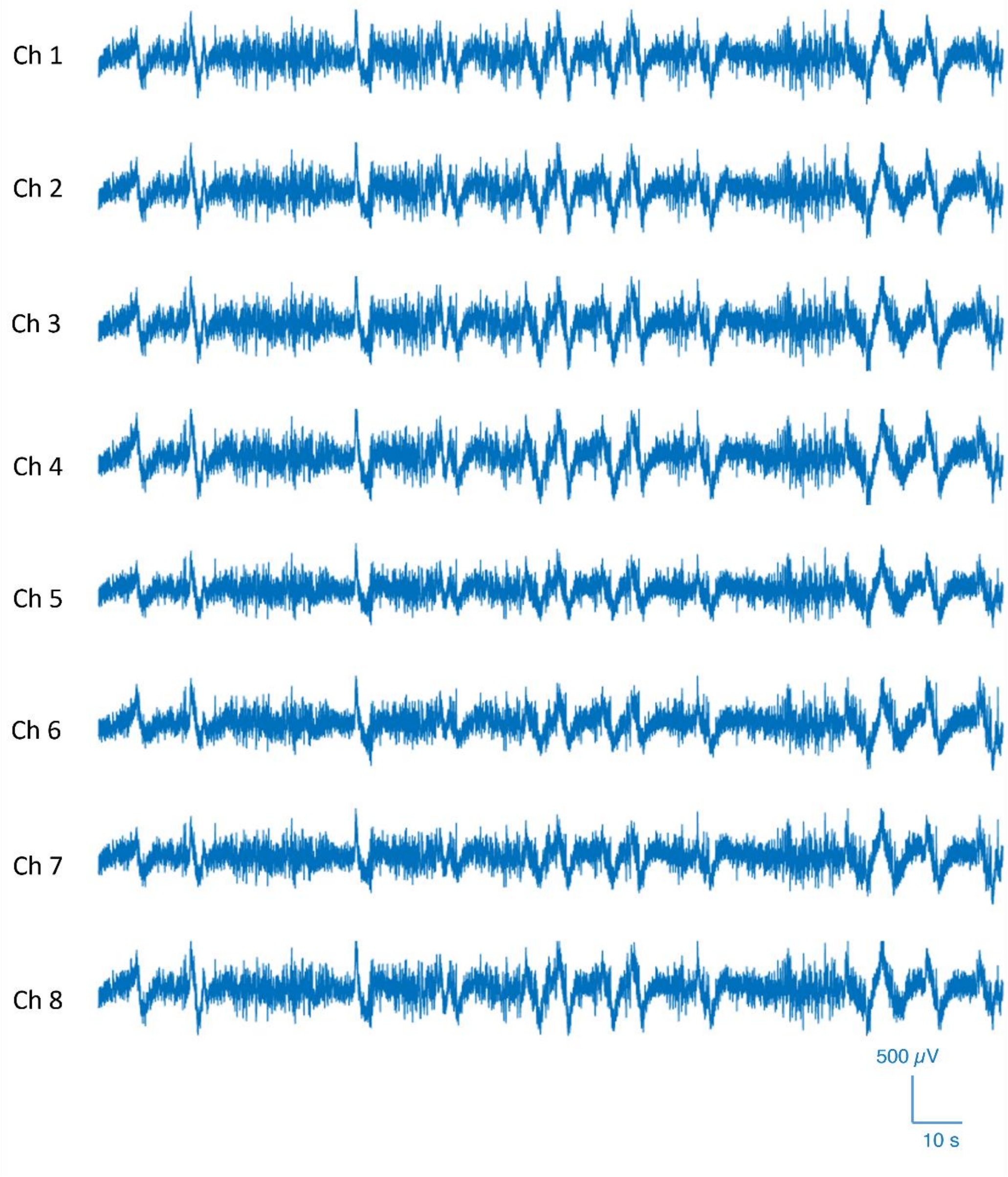


**Supplementary Figure 16. Full electrophysiology recordings by the mAxialtrode device.** Ch1 to Ch8 correspond to the EEG recordings from eight electrodes in the mAxialtrode.

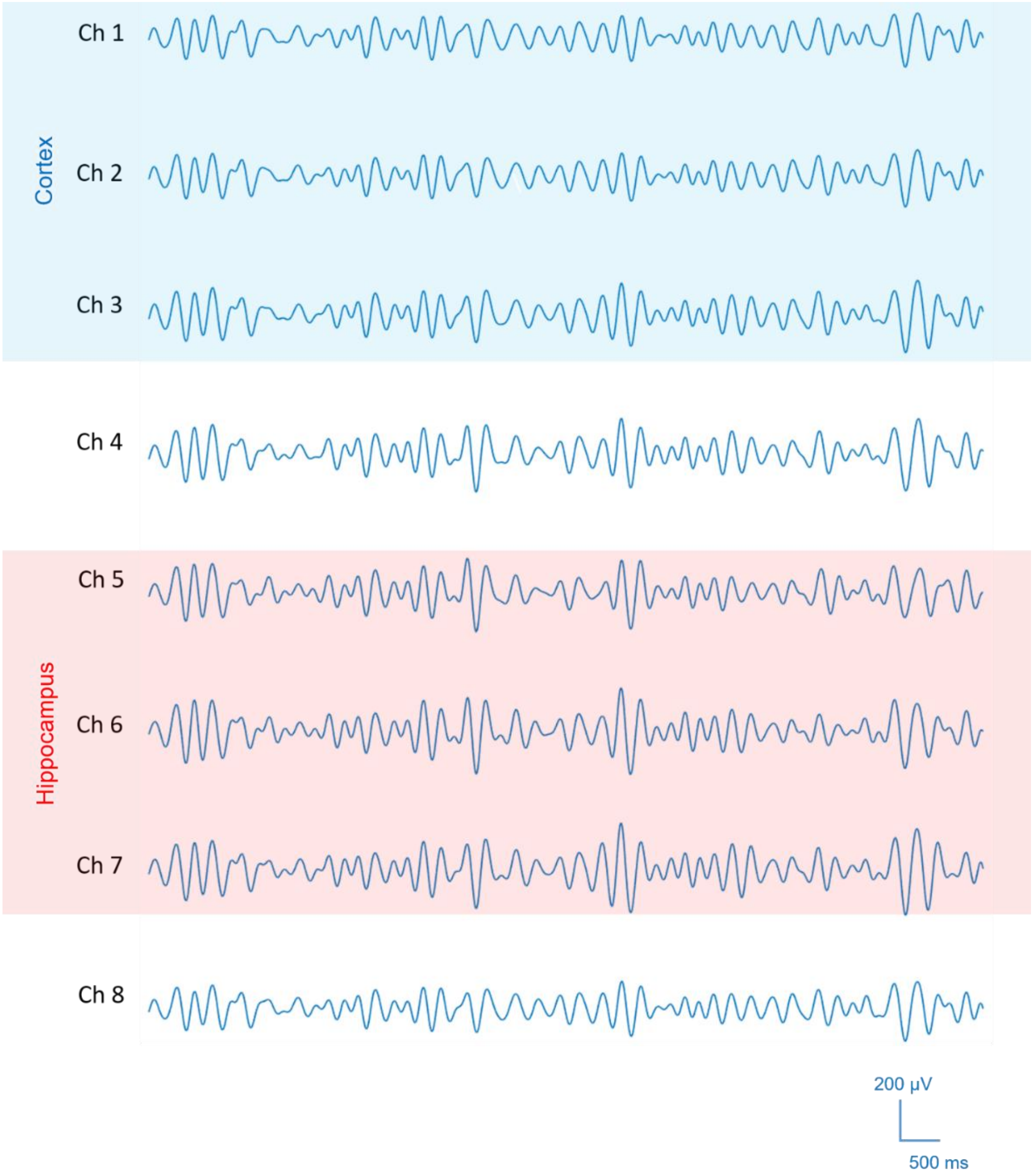


**Supplementary Figure 17. The theta rhythm component of the electrophysiology recordings by the mAxialtrode device.** The signals from Ch1, Ch2, and Ch3 correspond to the recording from the upper three electrodes positioned at the cortex (blue shadow) while the signals from Ch5, Ch6, and Ch7 correspond to the recording from the lower three electrodes that are aligned to the hippocampus (red shadow) of the mouse brain.

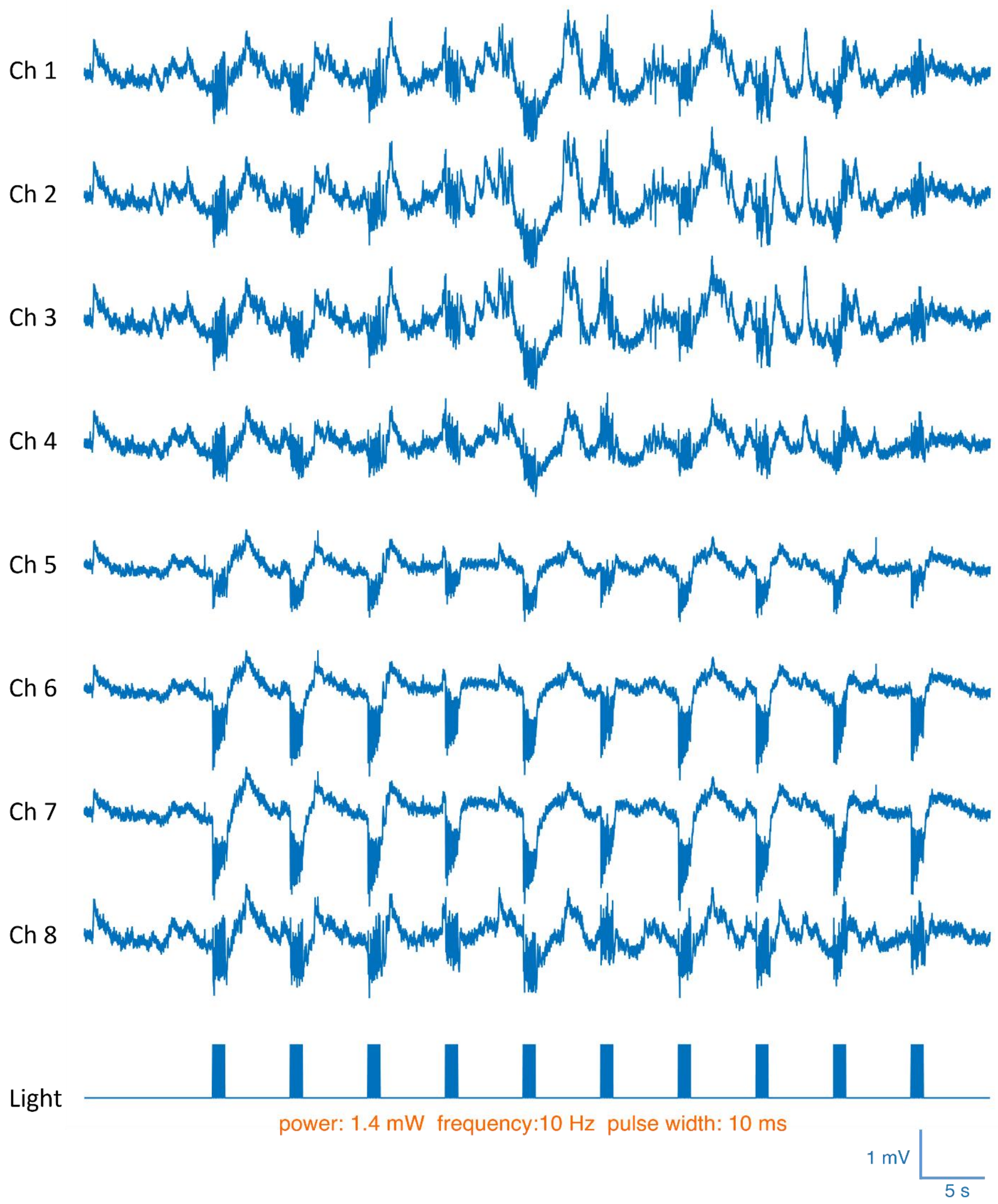


**Supplementary Figure 18. Typical electrophysiological recordings using the mAxialtrode device in optogenetic experiments demonstrate brain extracellular responses to light-induced activity**. This activity is elicited by 10 millisecond light pulses delivered at a frequency of 10 Hz and is collected by the eight integrated electrodes of the mAxialtrode.

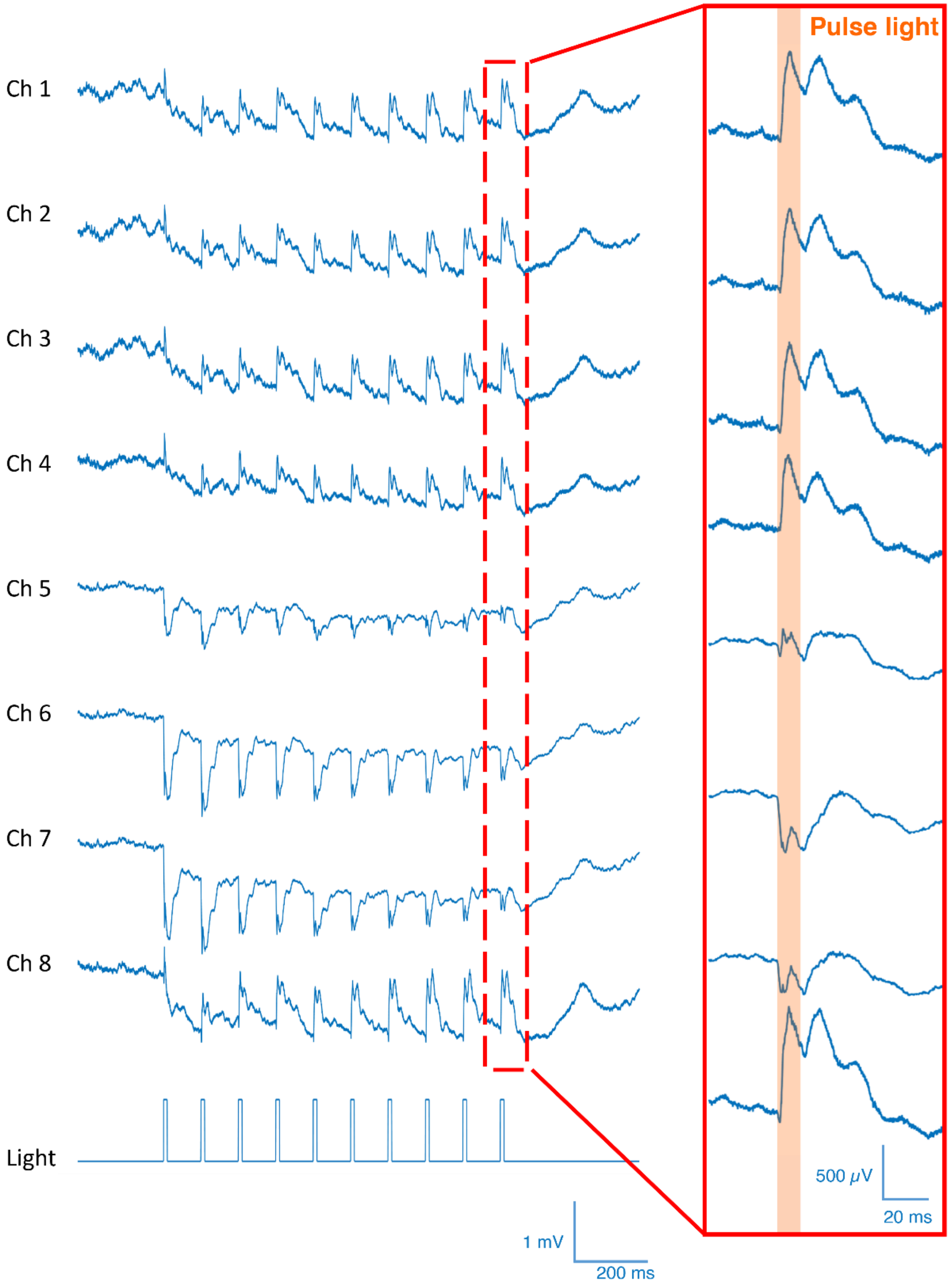


**Supplementary Figure 19. Typical 1s electrophysiological recordings in optogenetic experiments to show the details of the eight channels' traces in Supplementary Fig. 18.**

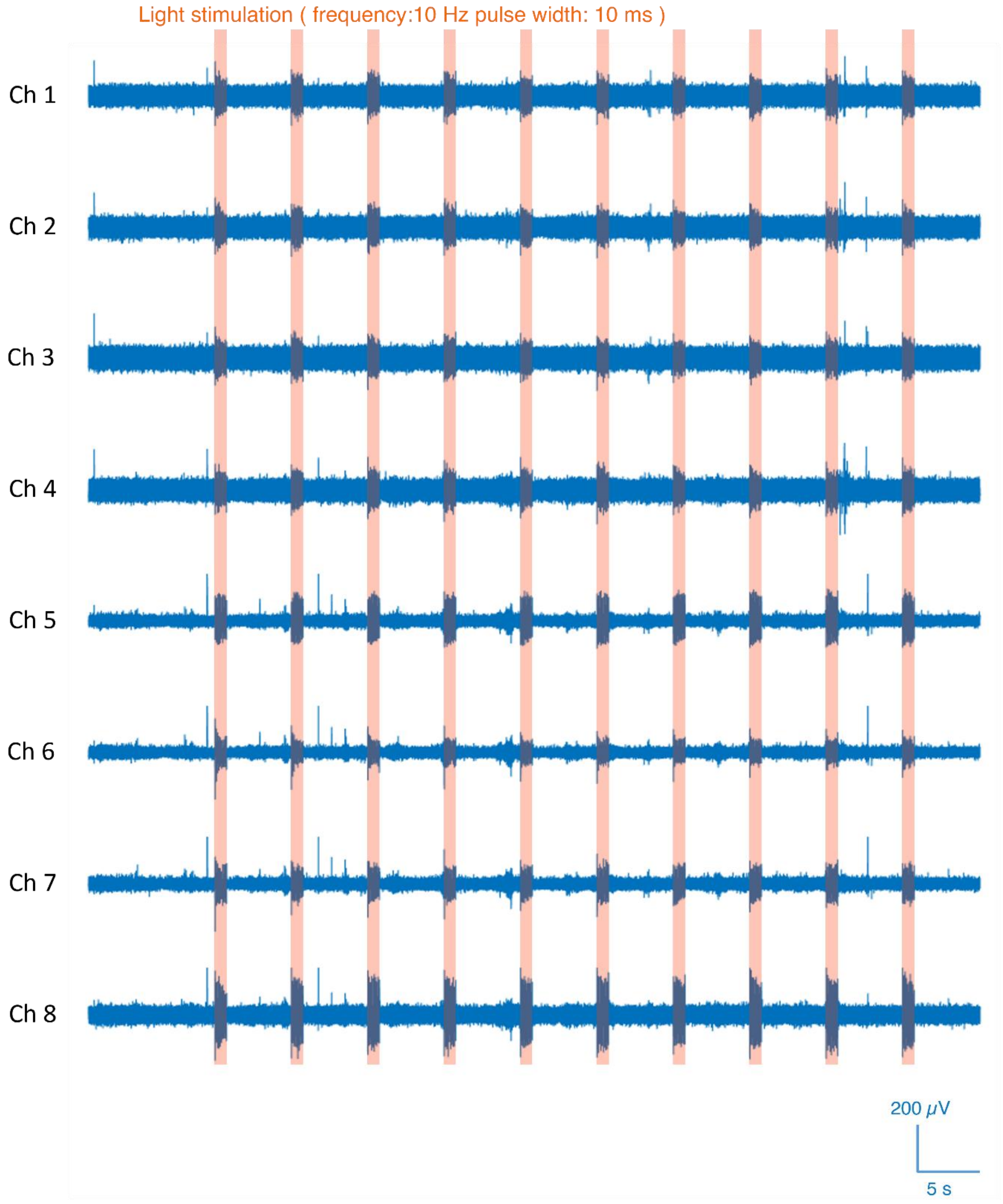


**Supplementary Figure 20. Optogenetic control of action potential firing in mouse brain.** For the recordings presented in Supplementary Fig. 18, a high-pass filter (>500 Hz) was used to highlight the recorded neural activities.

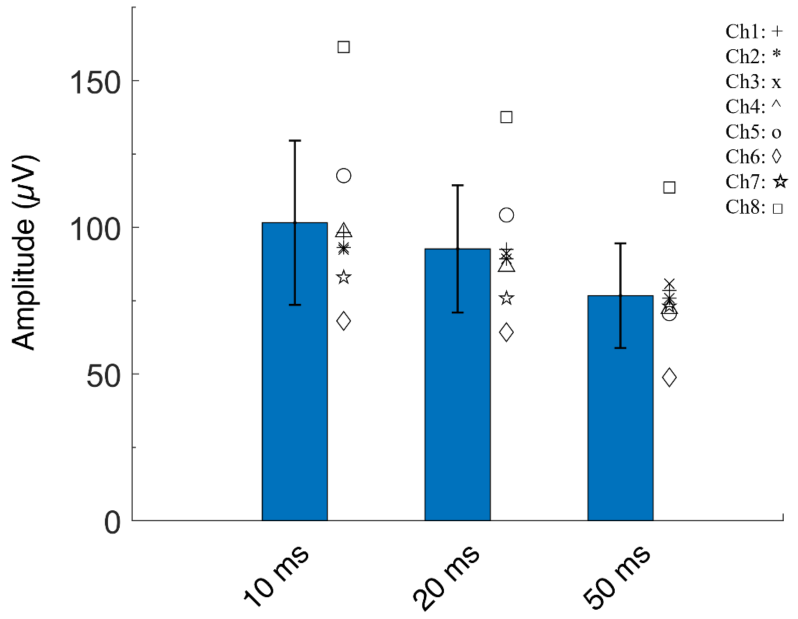


**Supplementary Figure 21. The evoked neural activity amplitude from all eight electrodes during optical stimulation with different pulse widths (10 ms, 20 ms, and 50 ms).**

| Print Settings Formlabs BioMed Clear | | | |
|---|---|---|---|
| Exposure Time | 13 s | Bottom Exposure Time | 26 s |
| Lift Distance | 8 mm | Layer Height | 30 $\mu m$ |
| Lift Speed | 60 mm/min | Retract Speed | 150 mm/min |
| Bottom Layer Count | 6 | Rest Time After Retract | 8,000 s |

**Supplementary Table 1. Print settings for Formlabs Biomed Clear resin on Phrozen Mighty 8K.**

| Pulse width (ms) | Spikes amplitude (μV) | Noise amplitude (μV) | SNR |
| --- | --- | --- | --- |
| 10 | 101.55 | 37.26 | 2.93 |
| 20 | 92.67 | 37.81 | 2.61 |
| 50 | 76.71 | 38.98 | 2.01 |

**Supplementary Table 2. The SNR of the spikes under different light pulse stimulation**
